# The origins and functional significance of bivalve genome diversity

**DOI:** 10.1101/2024.09.09.611967

**Authors:** Shikai Liu, Chenyu Shi, Chenguang Chen, Ying Tan, Yuan Tian, Daniel J. Macqueen, Qi Li

## Abstract

Bivalves are famed for exhibiting vast genetic diversity of poorly understood origins and functional significance. Within bivalves, oysters are an ancient group showing remarkable genetic and phenotypic variability alongside extensive adaptability, serving as an ideal system to understand the origins and functional significance of genomic diversity. Here, we reveal the divergent genomic landscape of *Crassostrea* oysters, characterized by a history of genome size reduction associated with transposable elements (TEs). By constructing a haplotype-resolved genome for Kumamoto oyster *C. sikamea*, we demonstrate the widespread presence of haplotype divergent sequences (HDS); genomic regions present in just one haplotype. Combined with population resequencing, we define the role of genomic divergence driven by TEs in shaping oyster genetic diversity. Comparisons of haplotype-resolved genomes across four bivalve orders reveal that while extensive HDS is common, its origins may differ markedly. We show that HDS are a hotspot of genetic innovation, harboring rapidly evolving genes of various evolutionary ages, while also strongly influencing gene expression phenotypes. A widespread lack of allele-specific expression shared among oyster individuals indicates that haplotype polymorphism provides a key source of expression variation, promoting phenotypic plasticity and adaptation. These findings advance understanding on the origins of genomic diversity and its role in adaptive evolution.

## Main Text

Genome diversity is the foundation of species evolution and shapes phenotypic diversity^1,2^. However, the mechanisms underlying genome diversity and its contribution to adaptive evolution remain insufficiently explored, particularly in species that maintain extensive genetic variation, such as bivalve molluscs, a highly diverse and geographically widespread group comprising over 9,000 species dating back to the Cambrian period^3^. The extensive presence-absence variation (PAV) observed among bivalve individuals and species underscores the remarkable genome diversity within this animal group^4^. PAV is an extreme form of structural variation, where certain genomic regions or genes are present in certain individuals yet absent in others^5^. Within an individual genome, PAV can appear as haplotype divergent sequences (HDS, i.e. regions unique to each haplotype) between homologous chromosomes, depending on the combination of haplotypes inherited from both parents. In the context of Mendelian inheritance, HDS may become dispersed or intermixed at the population level due to recombination and subsequent independent assortment of genetic material during meiosis. Therefore, characterization of PAV is needed to understand its role in adaptation at the population level^6^, while studying genome-wide haplotype differences can provide insights into the origins of genome diversity and its impact on genome functional features and gene expression.

Advancements in genome sequencing and assembly have greatly enhanced understanding of genetic variation in non-model species with complex genomes. Genomic features obscured in unphased genome assemblies can be revealed precisely using haplotype-resolved assemblies^7^, which are essential for understanding the basis for divergence among chromosomes (including HDS) and the significance of maintaining homomorphic or heteromorphic chromosomes^8^. Haplotype-resolved assemblies have recently been reported for domesticated plants with highly heterozygous genomes, providing rich insights into the genetic basis of heterosis and the evolutionary history of key crop species ^9–11^. Whole genome alignment^12^ has proven extremely useful to study HDS, specifically by identifying the unaligned segments between homologous chromosomes in haplotype-resolved genomes^13^.

Bivalve genomes exhibit extensive genetic variation, characterized by widespread diversity between and within individuals^4^. Both high reproductive capacity and genetic load are considered key to the highly heterozygous genomes of bivalves^14,15^. However, the relative contribution of mutation accumulation and gene flow to genome diversity remains poorly understood. Clarifying the origins of this diversity is important to understand how variation is maintained and shaped during evolution. Given its potential impact on gene expression^16,17^, understanding genomic diversity as a source of gene expression regulation may further provide important insights into the evolutionary success of bivalves.

Here, we explore the evolutionary origins of genome diversity in bivalves using haplotype-aware comparative genomics spanning different species and individuals from multiple orders, revealing the divergent genomic landscape of *Crassostrea* oysters. We uncover the origins of HDS and its role in shaping population characteristics by constructing a haplotype-resolved genome assembly for *C. sikamea*. By comparing haplotype-resolved genomes from four bivalve orders, we show that while extensive haplotype divergence is common to all, its composition and likely origin varies across species. We go on to establish the importance of oyster HDS as a source of new and rapidly evolving genes, before revealing its impacts on gene expression regulation using transcriptomics. We establish extensive variation in allele-specific expression among individuals, strongly coupled to haplotype polymorphism. Our findings advance understanding of genome evolution and gene expression variation in a non-model invertebrate clade showing extensive genetic diversity.

## RESULTS

### Genome reduction and divergence in *Crassostrea* species

Changes in genome content provide a key window into the evolutionary history of species^18^. Given the scale of genome diversity within and among bivalve species, large differences in genome size are expected to have arisen during evolution. To test this, we conducted a comprehensive comparison of genome sizes across diverse bivalve orders (Fig. 1a) using published data from the GoAT database^19^. A significant reduction in genome size was characteristic of species within the true oysters (Ostreida) (Fig. 1b), particularly among *Crassostrea* members. While bivalve genomes typically sum to around 1 Gb in length, *Crassostrea* species show much lower sizes (around 600 Mb) (Supplementary Data 1). Despite this, the number of annotated genes remains similar across phylogenetically diverse bivalve genomes, with comparable lengths of coding sequences (Fig. 1c). We observed extensive variation in gene family expansion and contraction across bivalves, reflecting high levels of genome diversity (Fig. 1c).

**Fig. 1.**
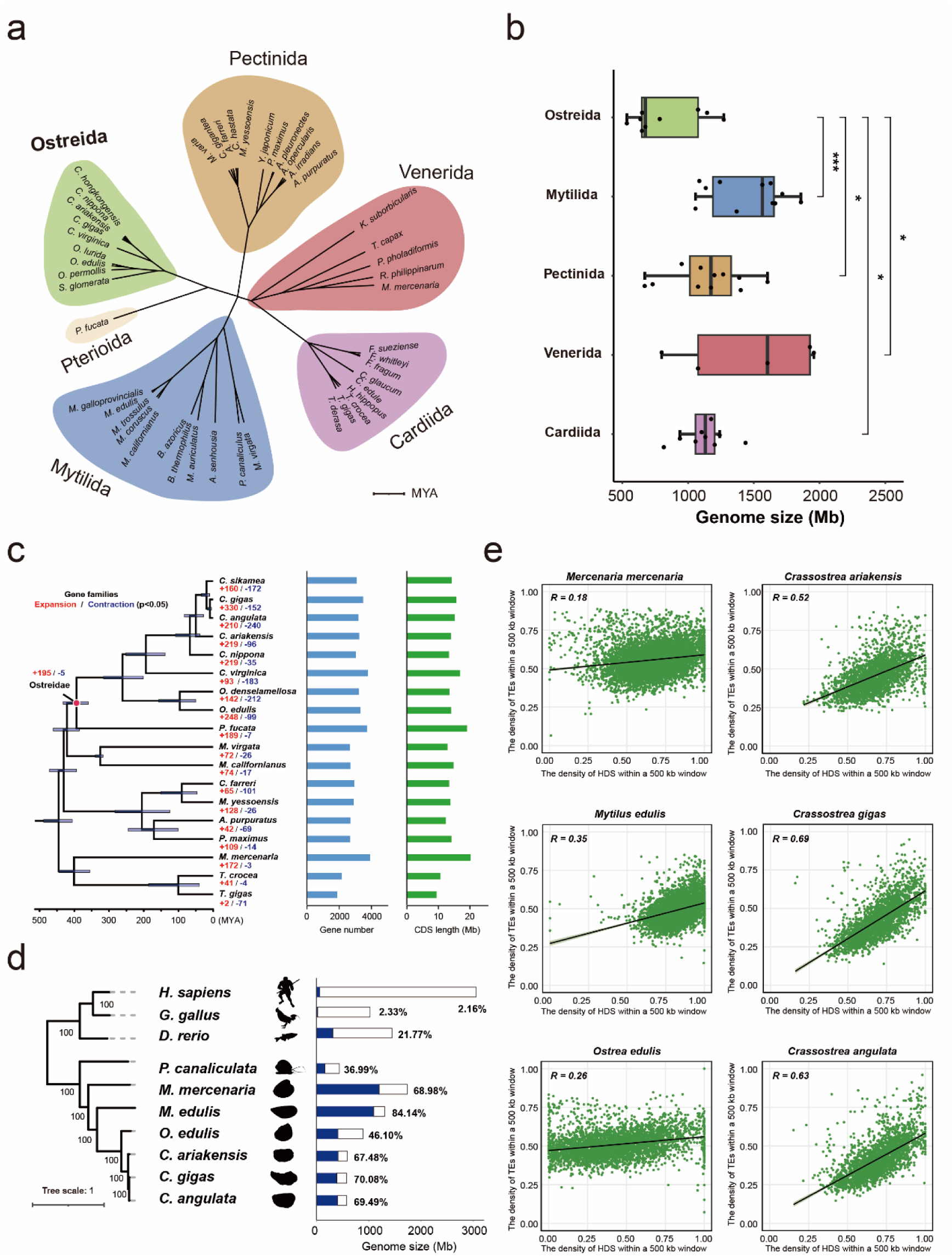
Comparative analysis of genome size, gene composition, and individual genomic variation across different bivalve genomes. **a** Phylogenetic tree of bivalve species with available genomic size data. Branch lengths were derived from the divergence times provided using TimeTree of Life^74^. **b** Comparison of species genome sizes across Pteriomorphia families (Mytilida, Ostreida, Petinida, Venerida, and Cardiida). P values were calculated using a two-sided unpaired Wilcoxon test; **c** Bivalve-wide analysis of gene gain and loss, indicating the number of genes gained (in red) and lost (in blue) for each species; counting gene number and total CDS length for each species. **d** Percentage of divergent sequences between individual genomes in representative species of bivalves and vertebrates. The bar chart shows the genome size for each species, with the blue segments representing the length of genomic differences, which are identified as divergent sequences that cannot be aligned within pairwise genome comparisons. Phylogenetic tree of species based on the alignment of single copy ortholog sequences. **e** Genome-wide distribution density comparison of divergent sequences and TEs for different Bivalvia species. The distribution density of each genomic component was calculated based on the proportion of sequences per 500 kb window.

To further understand the basis for genetic variation across bivalve species, we performed a series of pairwise genome comparisons (Supplementary Data 2). Despite potential inaccuracies due to differences in genetic background or assembly quality, our results illustrate extensive genomic differences across bivalve species (Fig. 1d). Notably, In the genome comparisons between individuals of several *Crassostrea* species, a similar proportion of significant genomic differences can be observed, characterized by a high proportion of divergent sequences that cannot be aligned with those of other genomes, indicating that extensive structural differences are widespread within this genus.

Transposable elements (TEs) are a recognized source of genome diversity in bivalves^4,20^. We found that genome divergence and TEs were significantly correlated in all bivalve species, but especially strongly for *Crassostrea* members (Fig. 1e), indicating that TEs have strongly shaped genomic variation in this genus.

### TEs shape haplotype diversity in oysters

To uncover the origins of genome diversity within *Crassostrea*, we assembled chromosome-level genomes for both haplotypes of a wild *C. sikamea* individual using HiFi and Hi-C data (Table S1, S2). The two genomes were 526.67 Mb (haplotype A) and 516.10 Mb (haplotype B) (hereafter: HapA and HapB, respectively) in length, both closely matching the estimated genome size by k-mer analysis (∼ 517.20 Mb, Fig. S1a). A series of gold-standard genome assembly assessment analyses showed both assemblies to be of excellent quality with high completeness and low error rates (Fig. S1b and S1c; Table S3). TEs comprise 41.29% and 39.59% of HapA and HapB, respectively (Supplementary Data 3). Interestingly, despite this similarity in global proportion, the contribution of different TE classes varied between HapA and B, especially for terminal inverted repeats (TIR) families (Fig. 2a). 30,052 and 29,609 protein-coding genes were annotated from HapA and HapB, which each showed high BUSCO completeness, while sharing extensive genome-wide collinearity (Table S2, Fig. S2a).

**Fig. 2.**
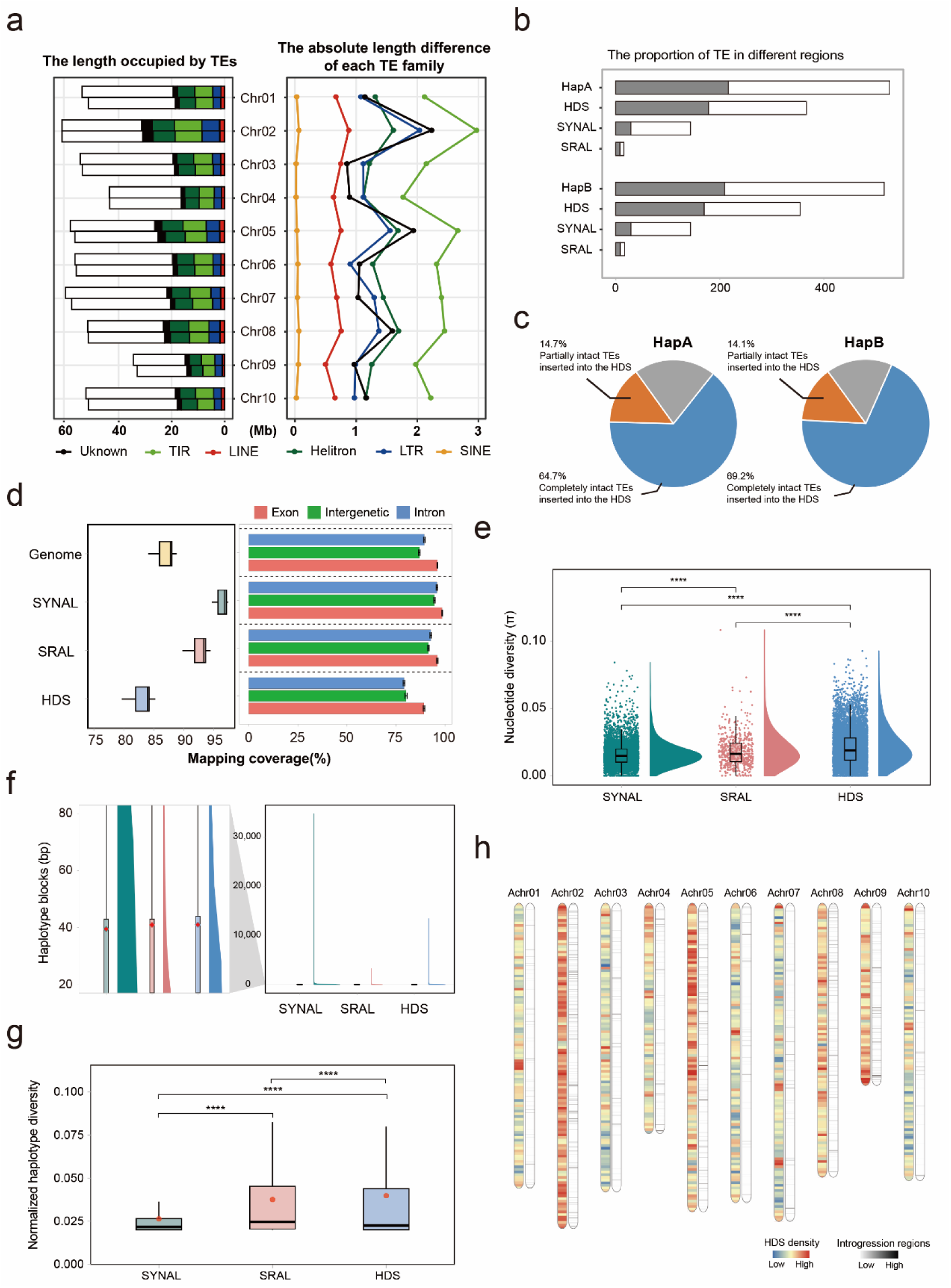
Transposable elements and genetic diversity within haplotype divergent sequences in *C. sikamea*. **a** The sequence length occupied by different classes of transposable elements (TEs) on each chromosome is shown on the left, with white bars representing chromosome length. The line graph on the right shows the absolute sequence length difference of each TE family within different TE classes between two haplotypes. **b** The proportion of TEs within syntenic region alignments (SYNAL), structural rearrangement alignments (SRAL), and haplotype divergent sequences (HDS) in the HapA and HapB genomes. **c** Proportion of intact TEs inserted in HDS. **d** Mapping coverage rate from population WGS data in different genome regions; **e** Nucleotide diversity of genes in different genome regions. **f** Length of haplotype blocks within different genome regions. **g** Normalized haplotype diversity within different genome regions. **h** Density distribution of HDS and population genetic introgression regions at the whole genome level.

Using HapA as reference, strikingly high proportions of haplotype divergent sequences (HDS, i.e. sequences that could not be aligned between haplotypes) were identified, constituting 69.59% and 68.69% of HapA and HapB, respectively (Table S4). The remaining sequences of the genome were divided into syntenic region alignments (SYNAL) and structural rearrangement alignments (SRAL), based on whether structural rearrangement occurs in the alignment (Fig. S2b). In total, 179.99 MB and 171.6 MB of TEs were inserted into the HDS of the HapA and HapB genomes, representing a significant portion of the whole-genome TEs (Fig. 2b). Specifically, 79.4% and 83.3% of intact TEs inserted entirely or partially within the HDS of HapA and HapB, respectively, which is more than in aligned/conserved parts of the genome (Hypergeometric test, p < 2.2e-16), indicating TE activity as a major driver for HDS (Fig. 2c).

Widespread HDS between haplotypes from a single *C. sikamea* individual suggests HDS will be evident at the population level. Using whole genome re-sequencing (WGS) with n=25 wild *C. sikamea* individuals, we found that fewer sequences align to HDS than conserved genomic regions (Fig. 2d). Moreover, nucleotide diversity of genes located within HDS regions was significantly higher than for aligned/conserved genomic regions (Fig. 2e). Although the length of haplotype blocks did not differ across genomic regions (Fig. 2f), haplotype polymorphism was significantly higher within HDS regions (Fig. 2g, p<0.01). To illuminate the impact of introgression on HDS and genetic diversity, we leveraged existing WGS data from three closely related species, including *C. ariakensis* (n=28), *C. angulata* (n=21), and *C. gigas* (n=21) (Fig. S3, Supplementary Note 1). We found evidence for some introgression between *C. gigas* and *C. sikamea* (Fig. S4). However, the degree of introgression was overall very limited (Table S5), with no correlation between candidate introgression regions and HDS (Fig. 2h). Therefore, we propose that genomic diversity within *Crassostrea* primarily originates primarily from TE proliferation, significantly affecting genetic diversity within extant populations.

### Comparative genomics of haplotypes across bivalves

To assess whether the extensive HDS observed in oysters is a general feature of bivalve evolution, we constructed haplotype-resolved genomes for a mussel (*Arcuatula senhousia*) and a scallop (*Mimachlamys varia*) (Table S6). Along with a published haplotype-resolved genome for pearl oyster (*Pinctada fucata*)^21^, high quality haplotype-resolved genomes representing four bivalve orders were compared. Extensive haplotype divergence was identified across all species compared (Fig. S5; Supplementary Data 4). The length-density distribution of HDS varied across bivalve groups, with a greater proportion of shorter HDS regions in oysters (Fig. 3a). A strong positive correlation was observed between the distribution of HDS and TEs in *C. sikamea* (R = 0.75), whereas this relationship was less pronounced for the three other bivalve orders (Fig. 3b).

**Fig. 3.**
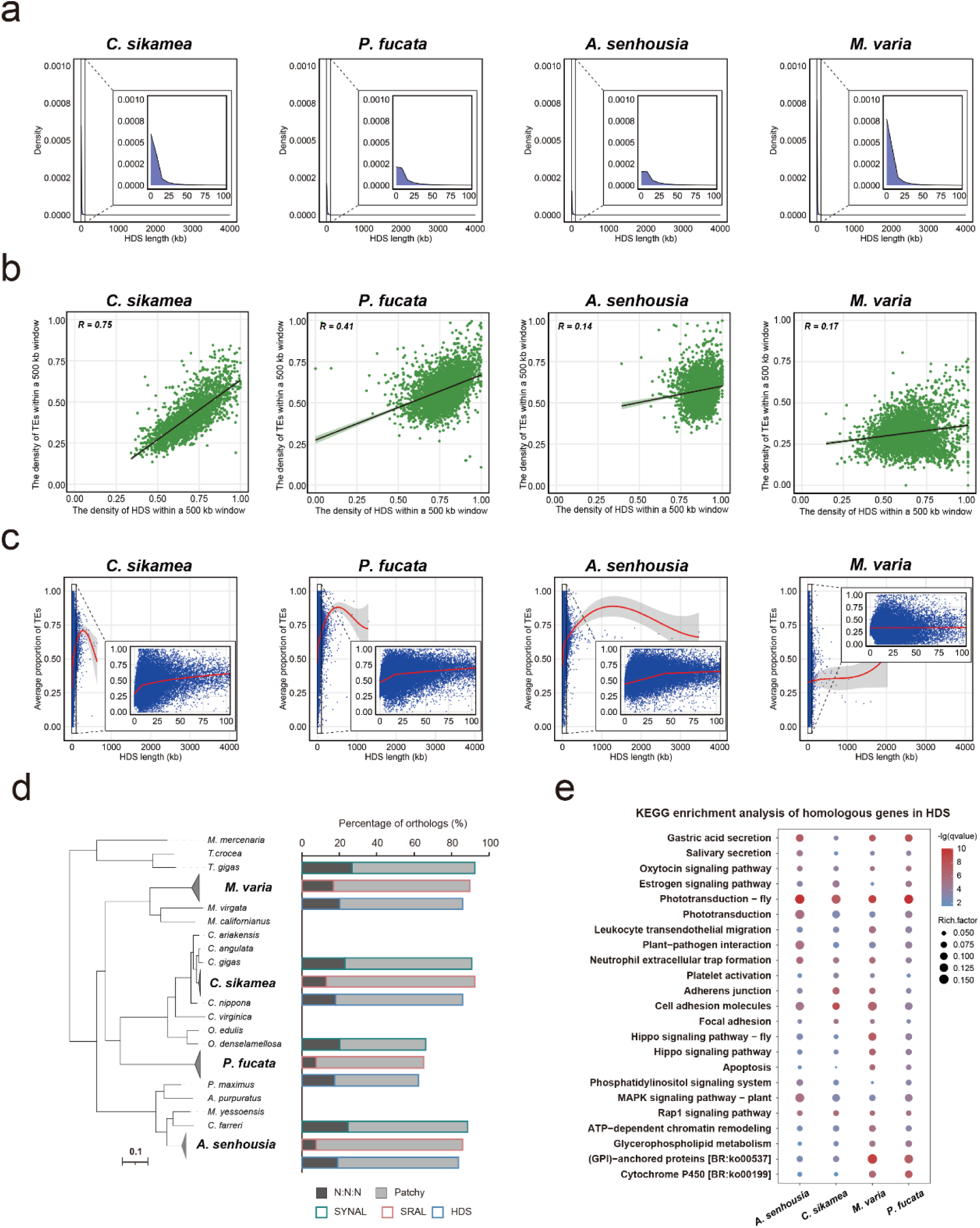
Comparative analysis of haplotype divergent sequences in bivalve genomes. **a** Density distribution of HDS lengths in four haplotype-resolved genome assemblies. **b** Density distribution comparison of HDS with TEs along the genome across each haplotype-resolved genome. The distribution density of each genomic component was calculated based on the proportion of sequences per 500 kb window; **c** Comparison of the average proportion of TEs in HDS of different lengths across each haplotype-resolved genome. **d** Distribution of homologous genes within syntenic region alignments (SYNAL), structural rearrangement alignments (SRAL), and haplotype divergent sequences (HDS) across the genomes of *C. sikamea*, *P. fucata*, *A. senhousia*, and *M. varia*. “N:N:N” indicates orthologous genes present in all species and “patchy” indicates the existence of other orthologs that are present in at least one genome. **e.** KEGG enrichment analysis of homologous genes within the HDS between haplotypes of *C. sikamea*, *P. fucata*, *A. senhousia*, and *M. varia*. Only the KEGG pathways that were significantly enriched in all four species were presented.

The distinct relationship between TEs and haplotype divergence across diverse bivalve species points to different mechanisms of genome evolution. To better understand this, we quantified the proportion of TEs within HDS across haplotypes for each species (Fig. 3c). In *C. sikamea*, *P. fucata*, and *A. senhousia*, TE content increases with HDS length, with evidence for an inflection point, where the rate of TE accumulation becomes limited. In *C. sikamea*, this inflection occurs earlier, and the initial TE content is lower, pointing towards a stronger inhibition mechanism for TEs in oysters. In contrast, TE content in *M. varia* does not change with increasing HDS length, suggesting factors beyond TE activity are shaping differences in haplotype divergence across different bivalve species.

We further identified a lower proportion of homologous genes within HDS compared to the genome-wide background for all species (Fig. 3d). Although the overall proportion of homologous genes within HDS was low, conserved genes within HDS were enriched in several functional pathways across all four species, including phototransduction, cell adhesion molecules, and rap1 signaling pathway (Fig. 3e). However, a larger number of biological functions were enriched either in a species-specific fashion, or in other pairwise combinations between species (Supplementary Data 5). This suggests that, while HDS may undergo rapid evolution, they harbor genes contributing to both conserved and lineage-specific biological functions.

### HDS as a source of young genes and adaptive capacity

To further understand the functional significance of haplotype divergence in bivalves, we explored the evolutionary and functional properties of genes located within HDS compared to other genomic regions. Gene clusters sharing high similarity (protein-level identity >90%; coverage >80%) were classified as recently duplicated genes in four bivalve genomes. We found these genes to be highly overrepresented within HDS in all cases (Fig. 4a; Hypergeometric test, p<2.2e-16), showing that HDS is a source of recently duplicated genes. Transcriptomic analysis from three tissues of *C. sikamea* revealed that genes located in HDS regions showed significantly lower median gene expression compared to the genome-wide background (Fig. S6), consistent with past findings that evolutionarily young genes often show low expression^22^, a hallmark of weak purifying selection.

**Fig. 4.**
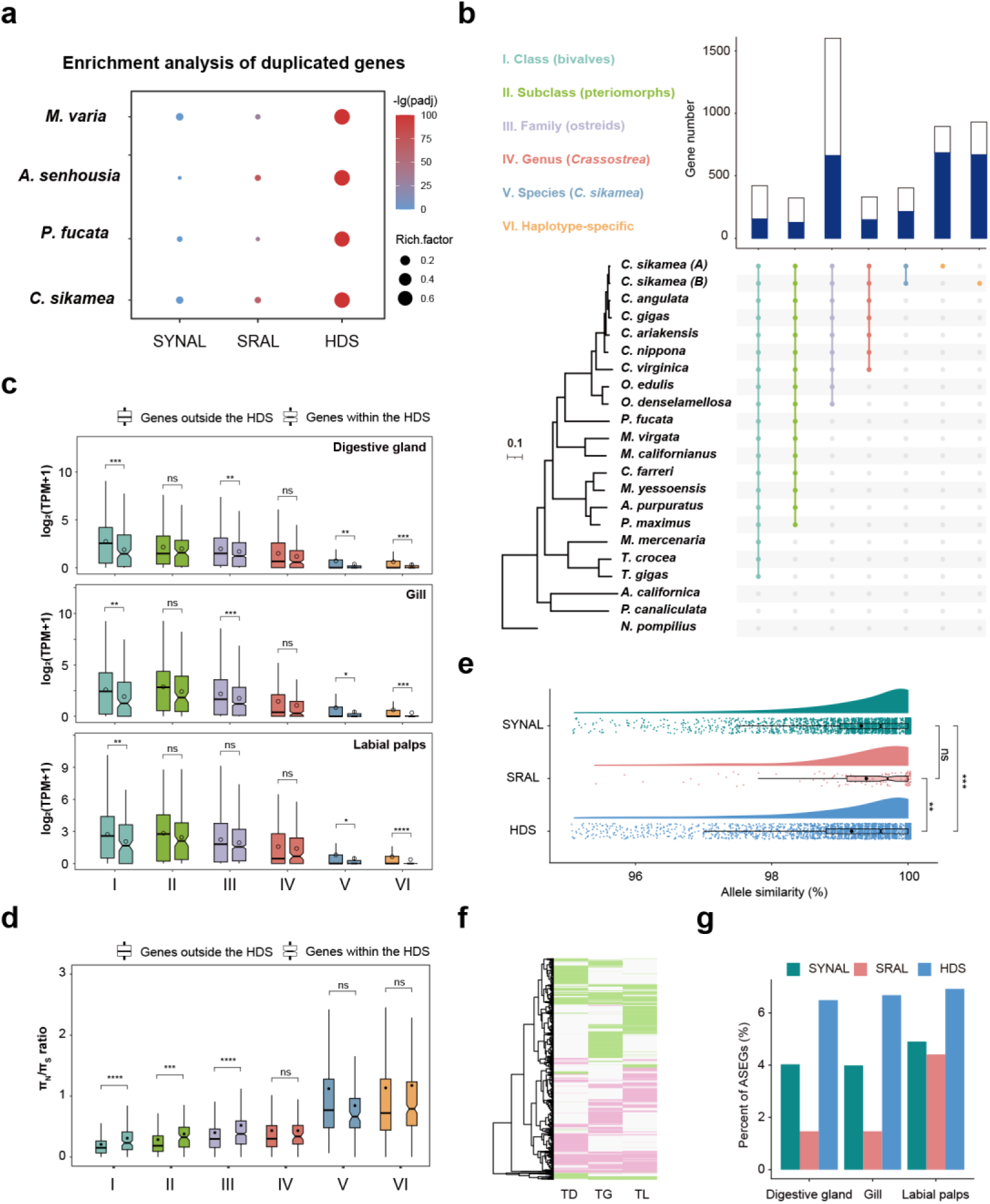
Impact of haplotype divergence on gene evolution and allelic expression. **a** Enrichment analysis of duplicated genes within syntenic region alignments (SYNAL), structural rearrangement alignments (SRAL), and haplotype divergent sequences (HDS) across the genomes of *C. sikamea*, *P. fucata*, *A. senhousia*, and *M. varia*. **b** Estimation of gene age according to conservation across different phylogenetic strata. The bar chart highlights the number of genes within the genome of *C. sikamea* that were identified as conserved across different phylogenetic clades, ranging from all bivalves through to genes found specifically in *C. sikamea*, including within HapA and HapB. The blue shading highlights the proportion of genes found within HDS. A phylogenetic tree of Mollusca is provided to highlight the relationship of species used in the analysis. **c** Summary of gene expression levels for genes categorized into different phylogenetic strata of the *C. sikamea* genome (data shown for digestive gland, gill, and labial palps). Significance testing was conducted using the Wilcox test. **d** The ratio of *π_N_* (nonsynonymous nucleotide diversity) to *π_S_* (synonymous nucleotide diversity) for genes categorized into different phylogenetic strata in the tested population of *C. sikamea*. **e** The proportion of allelic sequence similarity in different regions of the *C. sikamea* genome. **f** Allele expression patterns comparing digestive gland (TD), gill (TG), and labial palps (TL). **g** Summary of ASEGs across the genome vs. HDS regions comparing TD, TG and TL.

**Fig. 5.**
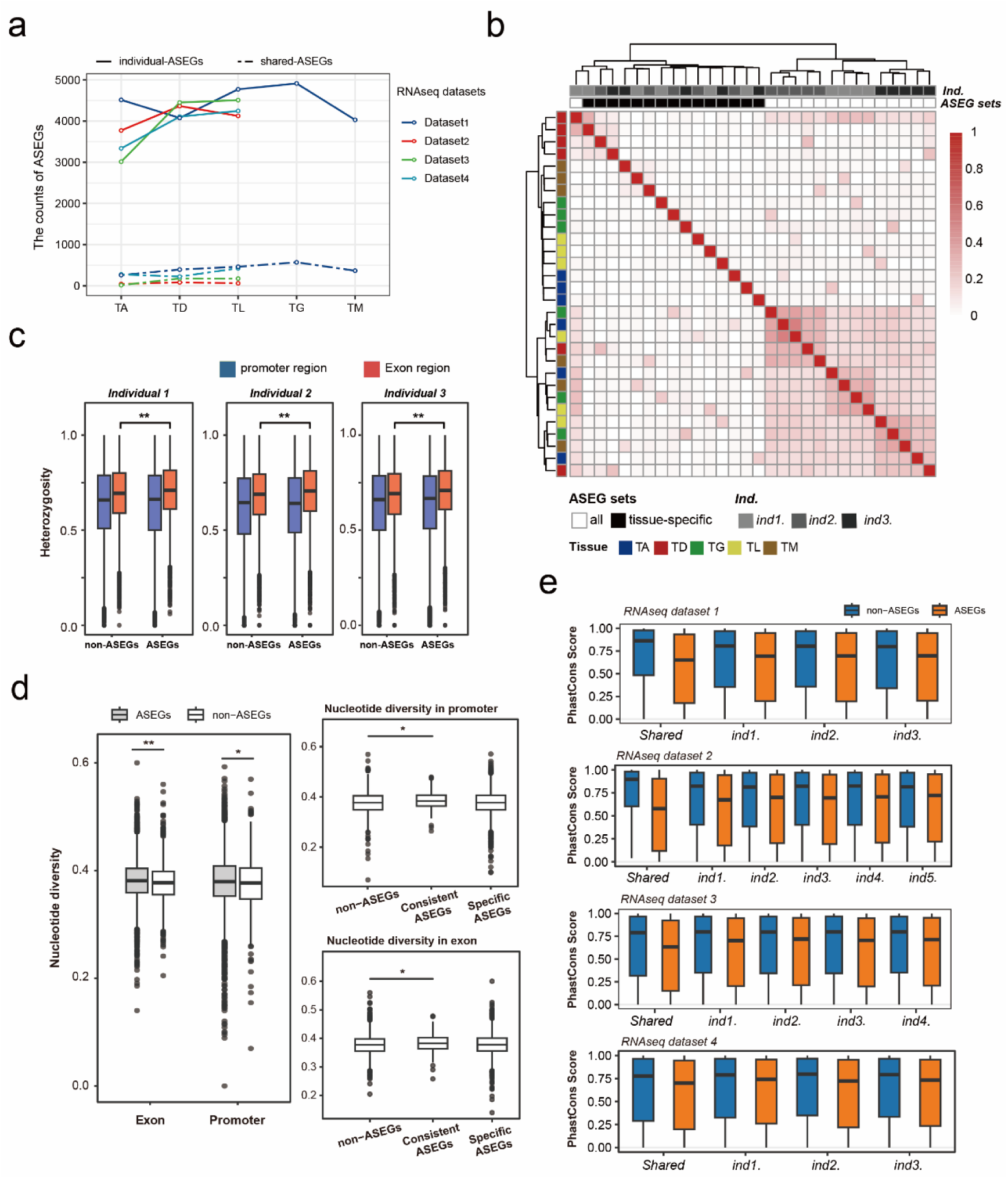
Expression profile, genetic variation and evolutionary constraint of allele-specific expression genes (ASEGs). **a** ASEG numbers from specific oyster individuals (individual-ASEGs, solid line) vs. all individuals (shared-ASEGs, dotted line) from various tissues (including TA, adductor muscles; TM, mantle; TD, digestive gland; TG, gill; and TL, labial palps) using each of the four RNA-seq datasets. **b** Heatmap representing similarity among the ASEG sets and tissue-specific ASEG sets from the RNA-seq datasets. Gene sets were clustered based on similarity represented by a cladogram on the top and left of the heatmap. **c** Heterozygosity of ASEGs and non-ASEGs within exon and promoter regions. **d** Nucleotide diversity of ASEGs and non-ASEGs among individuals within exon and promoter regions. Consistent-ASEGs indicate ASEGs shared among individuals that exhibit a consistent pattern of ASE across tissues; specific-ASEGs indicate ASEGs that are not shared among individuals. Statistics was conducted using the Wilcox test. **e** Evolutionary constraint of ASEGs and non-ASEGs. PhastCons scores of genes were determined across the genomes of nine Ostreidae species. ASEGs and non-ASEGs were identified for each individual using four RNA-seq datasets.

To more deeply explore the relationship between HDS and gene age, we split *C. sikamea* genes into different phylogenetic ‘strata’ spanning the breadth of bivalve evolution. We specifically binned genes into six groups based on their evolutionary origins being ancestral to: i) bivalves, ii) pteriomorphs, iii) ostreids, iv) *Crassostrea,* v) *C. sikamea* and vi) *C. sikamea-*specific genes restricted to one haplotype (Fig. 4b). These strata were defined based on orthogroups generated by OrthoFinder, which included genes that are unique within species from specific lineage, without considering orthologs from other species (Supplementary Data 6). Among all tested phylogenetic strata, a significant portion of genes are located within HDS (Table S7). Notably, the number of haplotype-specific genes located in HDS is more than expected by chance, as we detected 895 and 929 *C. sikamea-*specific genes unique to HapA and HapB, respectively, with 76.4% and 71.8% of these genes, respectively, located within HDS (Fig. 4b; Hypergeometric test, p < 2.2e-16). Haplotype-specific genes within HDS regions showed enrichment of KEGG terms including DNA replication, mitochondrial biogenesis, pattern recognition receptors, and NOD-like receptor signaling (Table S8).

In contrast to evolutionarily younger genes originating specifically during *C. sikamea* evolution (i.e. strata ‘v’ and ‘vi’), genes that arose at earlier stages of bivalve or oyster evolution (i.e. strata ‘i’ to ‘iv’) are comparatively more highly expressed in different oyster tissues regardless of their location within or outside HDS regions (Fig. 4c). The average expression levels of genes in strata ‘v’ and ‘vi’ was significantly lower in HDS regions in all tested tissues (Fig. 4c). A similar pattern was also observed for evolutionarily older genes in several tissues (strata ‘i’ to ‘iv’). Thus, HDS evidently harbors genes of diverse ages that show lower average expression compared to conserved genomic regions, supporting the idea that HDS genes are under more relaxed purifying selection.

We next asked if selective pressures acting on coding genes from the different phylogenetic strata were different between HDS and conserved genomic regions, comparing nonsynonymous (*πN*) versus synonymous (*πS*) nucleotide diversity (Fig. 4d). Consistent with our expression analysis, younger genes from strata ‘v’ and ‘vi’ show higher *πN* / *πS* ratios, indicating a faster evolutionary rate explained by weaker purifying selection and potentially positive selection. This pattern was similar for both HDS and non-HDS regions. In contrast, when genes from strata ‘i’, ‘ii’ and ‘iii’ are located within HDS, they exhibit significantly higher *πN* / *πS* ratios compared to genes of the same age located outside HDS. Hence HDS is associated with more rapid coding evolution for these evolutionarily older genes, which were enriched for GO terms related to the immune response and cellular homeostasis (Supplementary Data 7).

### Role of haplotype variation in gene expression evolution

Our new genome assembly allows us to further dissect the role of haplotype divergence on allelic variation and its role as a driver of gene expression evolution. Based on synteny and annotation of genes, 17,279 allele pairs were identified (57.92% of all annotated genes) (Supplementary Data 8), with an average nucleotide similarity of 99.2% in coding regions, and average Ka/Ks ratio of 0.16 (Fig. S7). In HDS, the proportion of identified alleles was 29.61%, which was lower than the whole genome level, while alleles in HDS regions exhibited significantly higher sequence divergence (Fig. 4e).

Using the identified alleles from HapA and HapB, RNA-seq data was used to explore allele expression. Direct quantification of allelic expression using the diploid genome provides an effective strategy to control for the effect of mapping bias (Fig. S8). As illustrated by an example in Fig. S9, we found evidence that alleles modified by nonsynonymous mutations or structural variations exhibited varying degrees of allele-specific expression (ASE) among tissues. Interestingly, varying degrees of ASE among tissues were observed (Fig. 4f) with the highest captured proportion of ASE in HDS, which is significantly higher than in SYANL or SRAL (Fig. 4g; Paired t-test, p <0.05).

### Extreme allele-specific expression linked to genome diversity

We next sought to expand the findings from our haplotype-resolved analysis of ASE, specifically to understand the significance of genome diversity in gene expression regulation in oyster populations. To achieve this, we developed a SNP detection method using (multi-tissue) RNA-seq data from different oyster individuals (Methods; Fig. S10). Our results demonstrate the reliability of SNP detection by RNA-seq (Supplementary Note 2; Table S9), allowing us investigate ASE across individuals and tissues to explore relationships with genome diversity. We first captured all genes showing ASE (ASE genes, ASEGs) in each tissue sample (Table S10). Among different tissues from the same individuals, a strong prevalence of tissue-specific ASEGs was identified, suggesting widespread tissue-specific *cis*-regulation. We observed similar expression of tissue-specific ASEGs across individuals, which was most pronounced in the digestive gland and mantle (Fig. S11).

To understand the functional significance of ASEGs, we compared four ASEG sets from each tissue of different oyster individuals (Fig. S12, Supplementary Data 9). Across each dataset, we observed numerous individual-specific ASEGs, while inter-individual (i.e., shared) tissue-conserved ASEGs were comparatively rare (Fig. 6a). Clustering analysis revealed that the similarity of ASEGs among individuals was significantly lower than within different tissues of the same individual (Fig. 6b). This suggests that ASEGs are more influenced by genetic background related to haplotype divergence than selective pressures associated with tissues during oyster evolution.

To unravel a potential evolutionary scenario for the occurrence of ASEGs, we investigated ASEG heterozygosity and found it significantly higher than for genes lacking ASE (non-ASEGs) in exon regions. However, heterozygosity in promoter regions lacked a direct correlation with ASE (Fig. 6c). To verify the connection between the extensive differences in ASEGs among individuals and haplotype polymorphism, we explored the relationship between genetic divergence among individuals and the occurrence of ASE. We found that ASEGs exhibit higher nucleotide polymorphism in both exons and promoters compared to non-ASEGs (Fig. 6d). Higher polymorphism may reflect neutral evolution or balancing selection, however, due to the small effect size, it is difficult to distinguish whether selection is strong or widespread.

The diversity of ASEGs captured among individuals is consistent with highly variable gene expression, pointing towards low evolutionary constraint on the affected genes. To test this, we obtained phastCons^23^ conservation scores using genome alignments spanning nine oyster species. Consistent with our hypothesis, we observe lower phastCons conservation scores for ASEGs specific to oyster individuals, compared to non-ASEGs. Moreover, with the increase in the number of tissues and individuals, non-ASEGs shared among individuals exhibited significantly higher conservation scores compared to ASEGs, which suggests ASEGs are less constrained by selection (Fig. 6e).

## Discussion

Understanding the origin and function of genome diversity is essential to deciphering the intricate relationship between genetic innovation and adaptive evolution. Based on our findings, we propose an evolutionary strategy in bivalves, where species enhance their adaptability by maintaining extensive individual-specific genomic diversity. A prime example is *Crassostrea*, a globally distributed genus that has adapted to the high-pressure environments of intertidal zones, maintaining a relatively small yet extremely diverse genome. Through the assembly of a haplotype-resolved genome for *C. sikamea* and precise analysis of haplotype divergence, we provide evidence that genome divergence shaped by transposable elements (TEs) enhances genetic diversity in oysters. Similar evolutionary mechanisms are present in selfing species, such as *C. elegans*^12^ and *A. thaliana*^24^, which implies this may be a common strategy for adaptive evolution.

Our comparisons of genomes representing four bivalve orders provided evidence that distinct mechanisms are creating and maintaining haplotype divergence and genome diversity, with notable differences in the contribution of TEs. For *Crassostrea* species, TE distribution within haplotype divergent sequences (HDS) follows a similar trend to species from the orders Mytilida and Pectinida. However, compared to other bivalves, there is a stronger correlation between TE content and HDS distribution across *Crassostrea* genomes, suggesting TE insertion may be more constrained. The high proportion of short HDS in *Crassostrea* genomes, along with limitations on TEs accumulation during HDS expansion, suggests that the expansion of HDS may be restricted, potentially linked to significantly reduced genome size in this oyster group. This association between genomic diversity and TEs in *Crassostrea* species positions them as a compelling model for studying the influence of TEs on genomic structure and diversity. For scallops, a significantly different composition of HDS compared to other tested species suggests other processes not driven by TEs may play important roles in bivalve genomic diversity. Constraint limiting the impact of TEs on scallop genome evolution may have contributed to the preservation of a conserved karyotype throughout their long history. The outstanding preservation of the ancestral bilaterian karyotype^25^, along with their hermaphroditic reproduction, could make scallops more dependent on genetic diversity arising from recombination, rather than TE activity. Notably, conserved homologous genes of diverse ages were identified within HDS across all investigated bivalve species, indicating a degree of conservation in the origins or functions of genomic diversity. This conservation likely contributes to species divergence in response to environmental pressures, emphasizing the intricate balance between evolutionary innovation and stability in bivalve genomes.

The potential function of haplotype divergence can be reflected in its impact on gene birth-and-death and expression regulation^26,27^. Our results demonstrate that HDS regions contain a significant proportion of the youngest duplicated genes, characteristic of rapidly evolving presence/absence variation (PAV) reported elsewhere^28^. This implies HDS represent hotspots for generating novel genetic diversity. Although our results suggest these recently duplicated genes are evolving rapidly under relaxed selective pressures, they likely serve as an important reservoir of genetic potential that can be used in adaptation in the future. Moreover, we identified many evolutionarily older genes within HDS regions showing comparatively high expression levels and accelerated evolution compared to genes of a similar phylogenetic age located outside HDS regions. These genes were enriched for immune and homeostasis functions, suggesting that conserved or older genetic elements within HDS could directly respond to selection pressures, supporting rapid adaptation within populations. Considering the extensive haplotype genome differences among bivalve species and the presence of conserved genes located within HDS, the high prevalence of genome divergence in these species may serve as a strategy that enhances their adaptive capacity.

In the context of genetic diversity, the different combination of divergent haplotypes can play a crucial role in shaping the complex genomic characteristics of individuals^12,29^. By utilizing transcriptomic datasets from various tissues, we evaluated the impact of genomic polymorphisms on expression regulation from multiple dimensions, providing evidence for a potential connection between complex genomic features and adaptive evolution. Our results strongly support extensive diversity of ASEGs in oyster populations. We found that the genotypes of conserved ASEGs exhibited more pronounced genetic polymorphism among individuals. Considering that oysters are classic profligate broadcast spawners that inhabit highly variable environments, including from generation to generation, we speculate that the maintenance of an extensive and diverse pool of ASEGs offers an adaptive evolutionary strategy, contributing to the high phenotypic plasticity observed among individuals.

Overall, this study offers strong evidence supporting the significance of genome haplotype divergence in promoting adaptation and evolutionary success, offering valuable insights into the evolutionary strategies employed by bivalves. These findings pave the way for future research to explore the broader implications of genomic variation and its roles in adaptive evolution using bivalves as a study system.

## Methods

### Sample collection and genome sequencing

Kumamoto oysters (*C. sikamea*) used for sequencing were collected from a wild population in Ningde (Fujian, China). Detailed sampling information is provided in Table S1. For population genomics, 25 oysters were sampled for whole genome resequencing (WGS). Adductor muscles were flash-frozen in liquid nitrogen and DNA extracted using a classic phenol-chloroform method. The concentration and quality of DNA was assessed using a Qubit fluorimeter (Thermo Fisher Scientific) and agarose gel electrophoresis. Sequencing libraries were constructed using the NEBNext Ultra DNA Library Prep Kit and sequenced on an HiSeq X Ten platform (Illumina) generating 150 bp paired-end reads.

For genome assembly, high molecular weight DNA was extracted from adductor muscle of one individual for single-molecule real-time (PacBio) and Hi-C sequencing. The PacBio library was prepared using a SMRTbell prep kit 3.0 and sequenced on the PacBio Sequel II System. The Hi-C library was prepared using the Dovetail™ Hi-C Library Preparation Kit and sequenced on the Illumina HiSeq X Ten platform generating 150 bp paired-end reads.

For genome annotation, transcriptome sequencing was conducted using various tissues from the same individual used for assembly. Total RNA was separately extracted from gill, digestive gland, and labial palps using a Trizol reagent (Invitrogen). RNA-seq libraries were constructed using the NEBNext^®^ Ultra™ RNA Library Prep Kit and sequenced on a NovaSeq 6000 platform generating 150 bp paired-end reads. Additionally, total RNA was extracted from tissues of multiple individuals, including mantle, gill, heart, digestive gland, labial palps, and adductor muscle. Equal amounts of RNA from these samples were pooled and subjected to full-length transcriptome sequencing. A PacBio Iso-Seq library was constructed using the SMRTbell prep kit 3.0 and sequenced on a PacBio Sequel II platform.

For gene expression analysis, adductor muscles, mantle, digestive gland, gill and labial palps were collected from three wild *C. sikamea* individuals. Tissue collection, storage, RNA extraction, and sequencing were as described above. Additionally, RNA-seq data was generated for *C. sikamea* at different growth stages from the same population. Samples were collected in March (n=4), June (n=3), and October (n=5) 2023. Adductor muscle, digestive gland and labial palps were collected and RNA-seq performed as described above.

### Haplotype genome assembly and comparison

Before genome assembly, Jellyfish was used to calculate k-mer frequencies (k = 19) before genome size, repetitiveness, and rate of heterozygosity were predicted with genomeScope2.0^30^. We assembled haplotype-resolved *C. sikamea* genomes using Hifiasm^31^. Juicer (v1.6)^32^ combined with 3D-DNA (v180419)^33^ was used for scaffolding. The same procedure was applied to assemble of haplotype-resolved genomes for *Mimachlamys varia*^34^ and *Arcuatula senhousia* using publicly data from the Darwin Tree of Life Project^35^.

Minimap2^36^ was used to align the two haplotype-resolved genomes using HapA as reference. The unaligned regions between haplotypes were classified as haplotype divergent sequences (HDS). SyRI^13^ was used to identify syntenic regions and structural rearrangements.

### Genome annotation of genes and repetitive elements

Genome annotation was performed separately for HapA and HapB using a comprehensive strategy combining evidence-based and *ab initio* gene predication. *De novo* identification and classification of repeats was performed using RepeatModeler^37^. RepeatMasker^38^ was used with a *de novo* repeat library for soft-masking. BRAKER3^39^ was used to annotate genes after soft-masking. The RNA-seq and Iso-seq data was supplied to the BRAKER pipeline along with OrthoDB v10^40^. For the genomes of several other representative species (i.e., *P. fucata*, *A. senhousia*, and *M. varia*) lacking RNA-seq data , we predicted coding genes using GALBA^41^, based on the annotations of closely related species. Functional annotation of protein-coding genes was carried out using EggNOG (v 5.0)^42^ and InterProScan (v 5.52–86.0)^43^. To achieve precise and comprehensive annotation of species-specific repeat libraries, we used a combination of structural and homology-based methods to annotate TEs with EDTA^44^. The genomes of several other representative species were also annotated for comparative analysis using the same approach.

### Population genomic analyses

WGS from *C. sikamea* individuals was used for variant calling. Fastp^45^ was used to trim adapters and low-quality reads with default parameters. Retained reads were aligned to the HapA genome using BWA^46^. Alignments were sorted and converted to BAM format using Samtools^47^. Duplicate marking and variant detection was performed using GATK4 following best practices^48^. Vcftools^49^ was used to filter SNPs with multiple alleles (>2) and missing rates >20% or minor allele frequencies < 5%. PIXY^50^ was used to calculate nucleotide diversity for each region. LDBlockShow^51^ was used to identify haplotype blocks. WhatsHap^52^ was used to generate phased haplotype sets for each individual. Subsequently, phased haplotype sets were imputed using Beagle^53^ for sites with a missing call rate > 20%. Following imputation, haplotype block diversity was calculated across genomic regions using Theta_D_H.Est^54^, normalized to the length of each region. SNPGenie^55^ was used to calculate nonsynonymous and synonymous nucleotide diversity.

We constructed an oyster SNP dataset incorporating our WGS data (n=25; *C. sikamea* individuals) with that for 70 individuals from three closely related species, including n=28 *C. ariakensis*, n=21 *C. angulata*, and n=21 *C. gigas* for population structure analysis. SNPs were identified as described above and pruned using Plink^56^ with the following parameters: *-indep-pairwise 50 5 0.2*, aiming to eliminate the influence of linkage disequilibrium (LD). Based on the pruned SNP dataset, a neighbor-joining phylogenetic tree was constructed in PHYLIP (version 3.695)^57^. Structure analysis was performed in Admixture^58^. *K* values were set from *K* = 2 to *K* = 9. The minimum CV error value appeared at K = 4. Principal component analysis was performed using Plink. For LD decay analysis, SNPs (retained post LD pruning) were analyzed using PopLDdecay^59^. The fixation index (*F*st) and nucleotide diversity (*π*) were calculated with a script (https://github.com/simonhmartin/genomics_genera). To assess the extent of shared DNA segments among individuals across the four species, identity-by-descent (IBD) block analysis was conducted using Beagle with the following parameters: *window = 100,000; overlap = 10,000; ibdtrim = 100; ibdlod = 5*.

We used the *D*-statistic^60^ (also known as the ABBA-BABA test) to capture candidate introgression loci among three ingroup taxa (*C. sikamea*, *C. angulata* and *C. gigas*) using *C. ariakensis* as an outgroup. Following detection of introgression among individuals at the genome level, we used a modified *f*-statistic (*fd*) to estimate the proportion of introgression sites at the population level. Using *C. ariakensis* as an outgroup, we expect equal counts of the two site patterns (ABBA and BABA) when incomplete lineage sorting results in inconsistent patterns at the sites. If the inconsistency is due to introgression, one of the site patterns is expected to be more prevalent. Leveraging the allele frequencies detected by GATK for each SNP, we used the R script calculate_abba_baba.r (https://github.com/palc/tutorials-1) to estimate the *D*-statistic, *fd* and *P*-value.

### Analyses of gene family, redundant genes, and expression profiling

A gene family cluster analysis was performed using annotated protein datasets of 21 representative molluscan species. To identify orthogroups and orthologs from the datasets, OrthoFinder (v2.5.4)^61^ was used including DIAMOND and OrthoMCL to call orthogroups based on sequence identity. The detailed procedure and subsequent phylogenetic analyses are described in Supplementary Note 3. Orthogroups present within specific phylogenetic strata were generated by OrthoFinder. These groups contain genes from the target species exclusively, without including genes from other compared species. To identify high similarity genes in the *C. sikamea* genome, we used CD-HIT^62^ with a sequence identity threshold of 90% and a minimum alignment coverage of 80%. For gene expression analysis, RNA-seq reads were mapped to the reference genome using STAR^63^, and expression levels quantified using StringTie^64^.

### Identification and comparison of alleles between haplotype genomes

Similar to previous reports^10,65^, *C. sikamea* HapA and HapB alleles were identified by assessing gene synteny, sequence similarity, and functional annotation results. Comparison of gene synteny between HapA and HapB was performed with WGDI^66^. Paired genes within each synteny block with the best BLAST score were considered putative alleles. These putative alleles were further filtered based on functional annotation results obtained using EggNOG (v5.0)^42^ and protein sequence length, retaining only those with identical functional annotations and sequence length differences < 25%. EMBOSS needle^67^ was used for pairwise comparison of allelic genes with default parameters. Allele sequence similarity score was calculated as the number of unsubstituted bases divided by the length of the alignment block. Finally, alleles with protein sequences exhibiting more than 5% divergence were filtered. Ka (number of nonsynonymous substitutions per non-synonymous site) and Ks (number of synonymous substitutions per synonymous site) values for allele protein-coding sequences were calculated by WGDI. For allele expression analysis, the RNA-seq data from the genome-sequenced individual was mapped to the diploid genome using STAR^63^, and expression levels quantified with StringTie^64^.

### Allele-specific expression (ASE) analysis

SNPs were detected using RNA-seq data to differentiate expressed alleles between haplotype genes using the GATK pipeline. Briefly, RNA-seq reads from the various sampled tissues of the same individual were combined to gain power in SNP detection, which were subject to stringent trimming with fastp. STAR was used with default parameters for read mapping on *C. sikamea* HapA. Uniquely mapped reads were then processed following GATK best practices for RNA-seq data to detect SNPs, which were filtered using “VariantFiltration” with two of the three suggested filters, “QD < 2” and “FS > 30”. Finally, we selected the SNPs with genotypes associated with each individual that Genotype Call Rate (CR) ≥ 20%, supported by at least 10 reads, and CR of all samples ≥ 50%^68^.

These SNPs were used to generate a pseudo-genome per individual by the “FastaAlternateReferenceMaker” tools from GATK. Each tissue sample sequence was aligned to the individual genome and then used to detect allele expression among each sample by phASER^69^. To assess ASE in each sample, we screened for read number imbalance between the alleles using a binomial test (binom.test R function). P-values were corrected using the Benjamini-Hochberg method with a false discovery rate of 0.05. The alleles with p-adj > 0.1 were considered to have no occurrence of ASE. We also used WGS data from each sampled *C. sikamea* individual for SNP detection and comparative analysis. SNP calling was done as described above. HiBLUP^70^ and PIXY^50^ were used to calculate heterozygosity and nucleotide diversity across different genomic regions, respectively.

### Detecting evolutionary constraint

Cactus^71^ was used to align the genomes of nine oyster species: *C. sikamea*, *C. angulata*, *C. gigas*, *C. nippona*, *C. hongkongensis*, *C. ariakensis*, *C. virginica*, *O. denselamellosa*, and *O. edulis*. Using the *C. sikamea* coding gene annotation, we first extracted 4-fold degenerate sites using hal4dExtract and used phyloFit^72^ to estimate an initial neutral model. Once the neutral model was calculated, using the Cactus alignment, we estimated *rho* (the expected substitution rate of conserved elements relative to neutrality) and calculated PhastCons scores for every site in the genome by phastCons^73^. We averaged the phastCons scores across the nucleotide sequences for each protein-coding gene to obtain the phastCons scores in gene-level. The phastCons scores range from 0 to 1 and represent the probability that a nucleotide sequence is under negative selection.

## Acknowledgements

This work was supported by the grants from the National Natural Science Foundation of China (32341060 and 42276112), the Key Research and Development Program of Shandong Province (2021ZLGX03), the National Key Research and Development Program of China (2022YFD2400300), and the earmarked fund for the Agriculture Research System of China (CARS-49). We acknowledge the support of the High-Performance Biological Supercomputing Center at the Ocean University of China. DJM was supported by Institute Strategic Programme funding to the Roslin Institute from the Biotechnology and Biological Sciences Research Council (awards: BBS/E/D/10002070 and BBS/E/RL/230001B).

## Author contributions

QL and SL initiated this project and obtained funding for the studies, SL conceived the project, coordinated and supervised the entire project. YT1 (Ying Tan) contributed to the sample collection and genome sequencing. CS and YT2 (Yuan Tian) assembled the genome of *C. sikamea*. CS, SL, DJM, and CC analyzed the data, CS and SL wrote the manuscript. SL and DJM revised the manuscript. All authors reviewed and edited the manuscript.

## Supplementary information

**Supplementary Information**

Supplementary Figs. 1–14, Tables 1–10, Notes 1–3 and references.

**Supplementary Data 1-10**

### Data availability

Sequencing data are available from NCBI SRA under BioProject accessions PRJNA1090262, PRJNA1090167, PRJNA1089862; The genome assembly and annotation data of the accessions is available at Figshare 10.6084/m9.figshare.26953708.

### Code availability

Custom code used for the analysis is available at https://github.com/Scy-bio/paper-anlysis-pipeline.

