## Supplementary Figures 1-14, Tables 1-10, and Notes 1-3 for "The origins and functional significance of bivalve genome diversity"

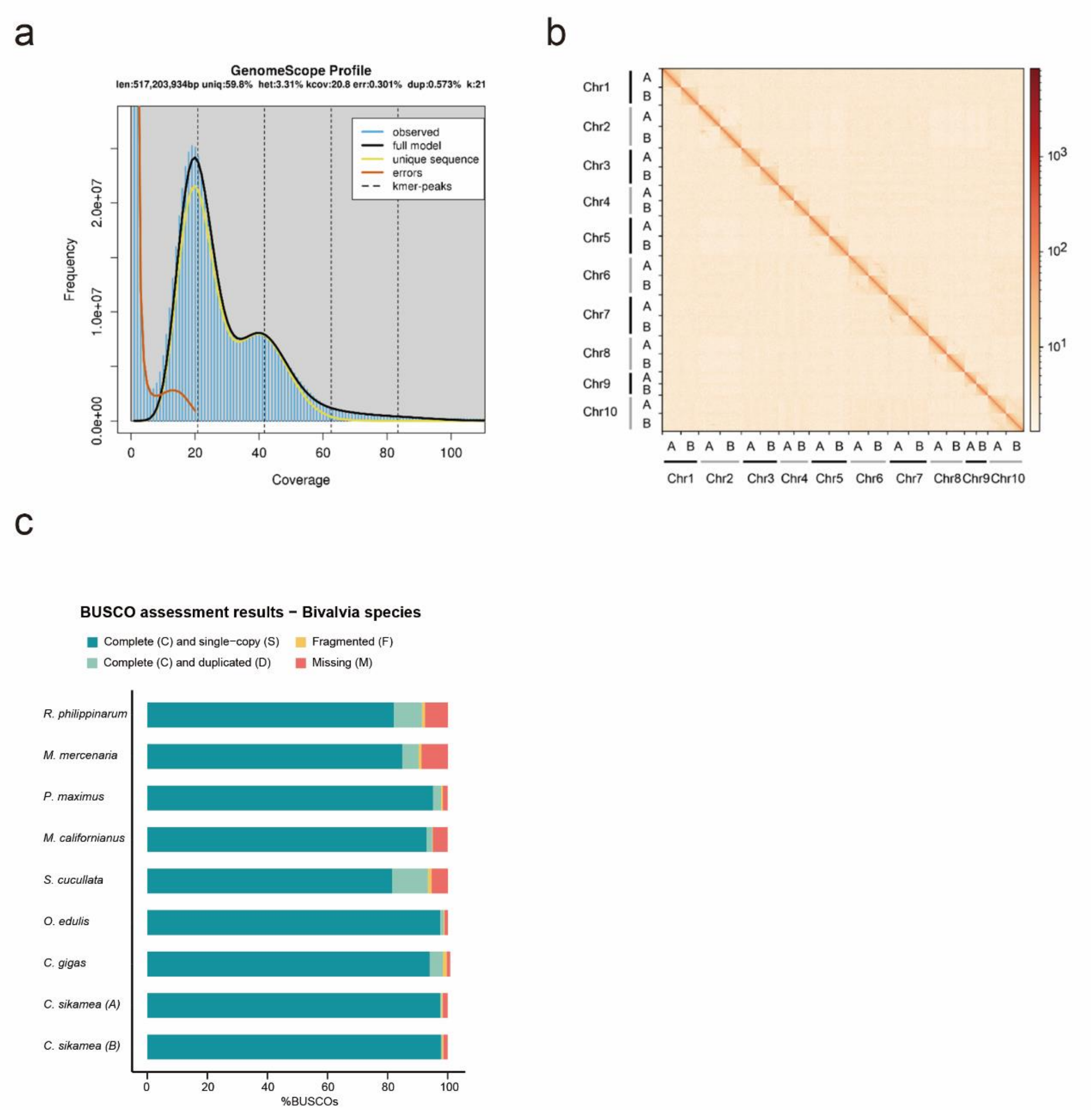


**Supplementary Figure 1. The assessment of *C.sikamea* genome size and of assembly completeness. a** K-mer spectra assessment for the genome assembly. **b** Hi-C interaction matrix of the *C.sikamea* diploid genome. **c** Genome completeness, as indicated by mollusca BUSCO repertoire (mollusca_odb10, n= 5,295), in genome assemblies of different bivalvia species.

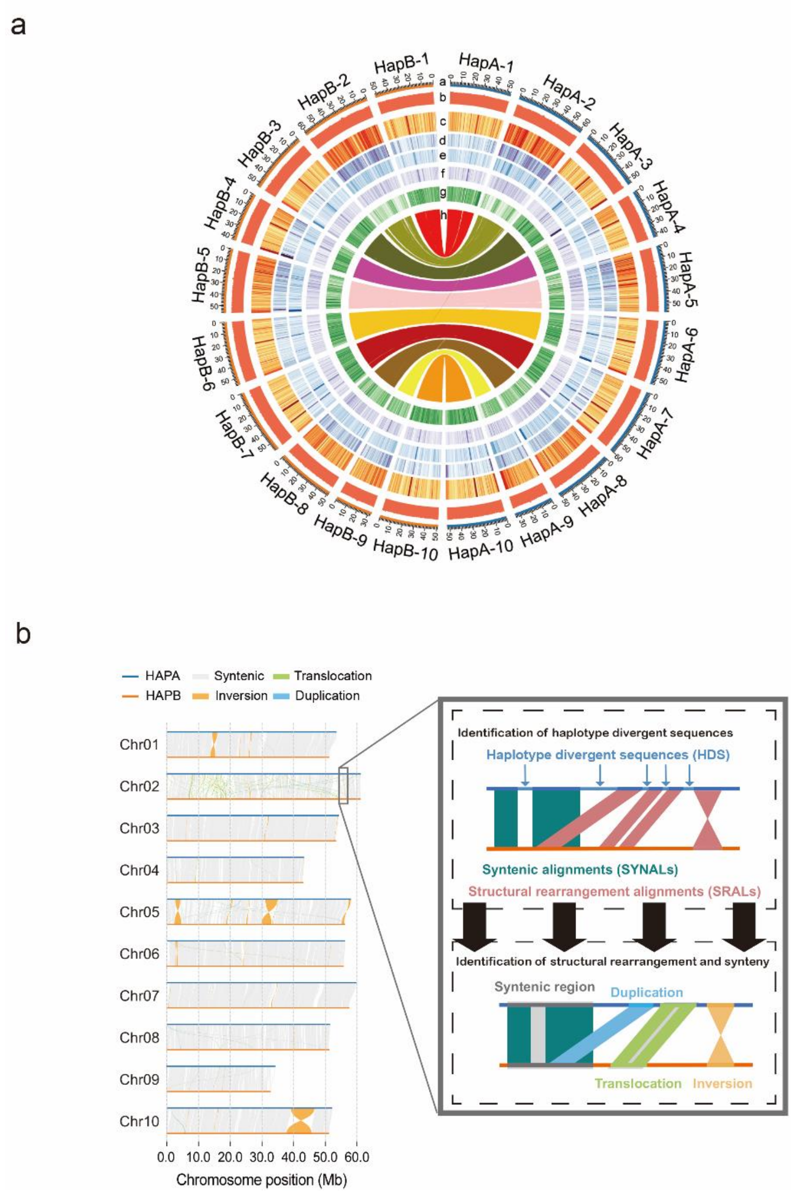


**Supplementary Figure 2. The assembly and comparison of *C.sikamea* haplotype-resolved genome. a** Circos plots showing 20 chromosomes (a), the distribution of GC content (b), transposable elements (c), LTR (d), TIR (e), Helitron (f), Gene (g), and the synteny of Gene. **b** Synteny and structural rearrangements are represented by colored lines connecting regions of the haplotype-resolved chromosome-level genome of *C. sikamea*. In the close-up view, the schematic diagram illustrates the definition of structural rearrangement and synteny region. The unaligned regions between haplotype were classified as haplotype divergent sequences (HDS) based on the positional relationship of annotation blocks. The remaining sequences of the genome are divided into syntenic region alignments (SYNAL) and structural rearrangement alignments (SRAL) based on whether structural rearrangement occurs in the alignment.

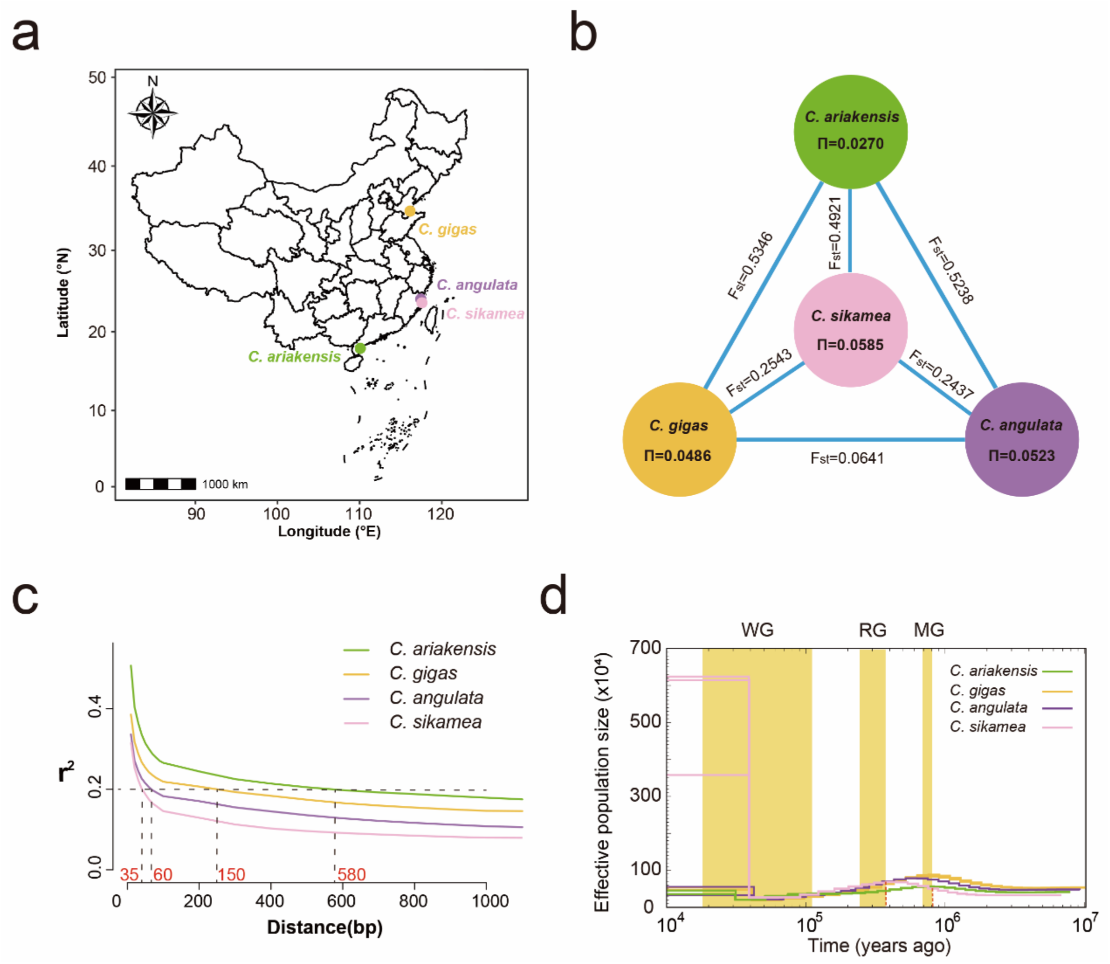


**Supplementary Figure 3. The population structure and demographic of four *Crassostrea* species. a** Sampling sites of *C. gigas*, *C. angulata*, *C. sikamea*, and *C. ariakensis*. **b** Nucleotide diversity (Π) and genetic divergence (*F*st, between sampled individuals) among four oyster species. **c** Decay of linkage disequilibrium in four oyster species. **d** The demographic history of the four oyster species was inferred using the pairwise sequentially Markovian coalescent (PSMC) model. Periods of the Mindel glaciation (MG, 0.68-0.80 million years ago), Riss glaciation (MG, 0.24-0.37 million years ago), and Würm glaciation (WG, 10,000-120,000 years ago) were indicated by yellow shading.

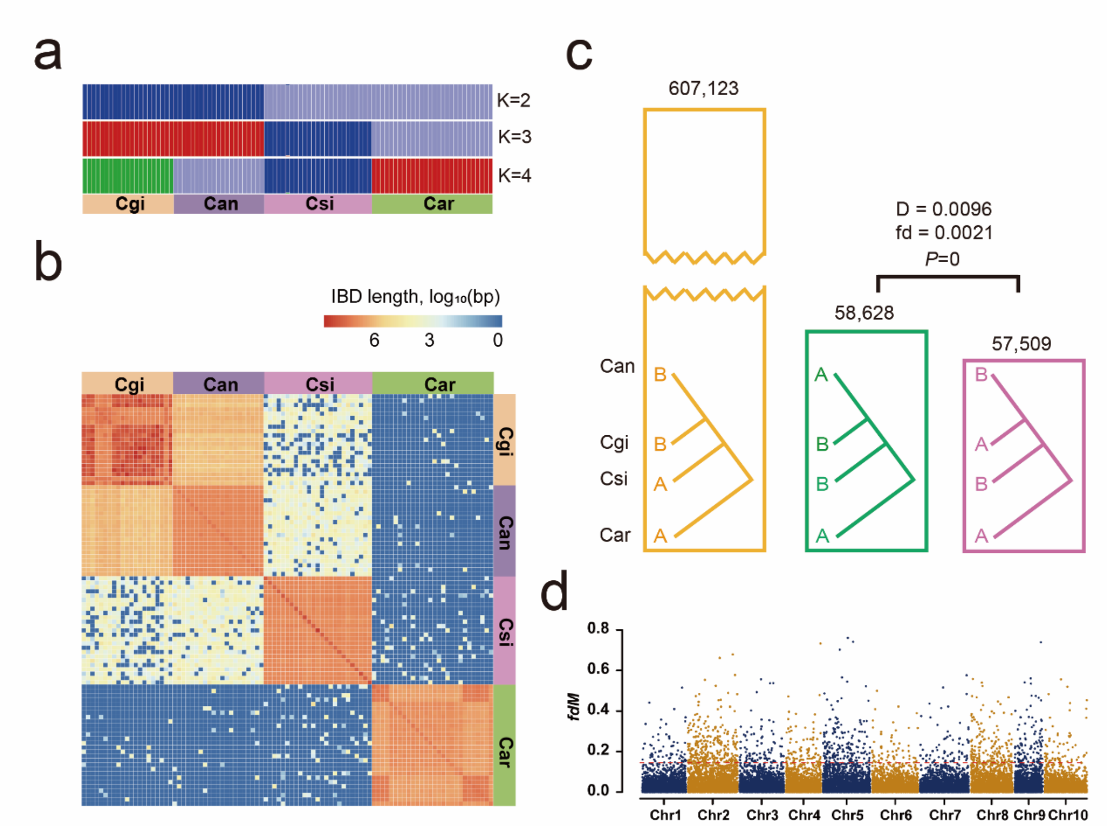


**Supplementary Figure 4. The introgression of four *Crassostrea* species. a** Admixture analysis was performed with K=4, where the CV error value was the lowest. The Admixture plot shows the inferred genetic components for each individual. The colors in the Structure plot represent the different genetic components present in each individual sample; *C. ariakensis* (Car), *C. sikamea* (Csi), *C. angulata* (Can) and *C. gigas* (Cgi)**. b** Estimated haplotype sharing between individuals of the four oyster species. The colors in the heatmap represent the length of identifcal by decent (IBD) blocks shared between individuals. **c** ABBA-BABA analysis of the four oyster species. The number of sites supporting different tree structures was shown in a bar plot. To test the null hypothesis of symmetry in genetic relationships, *D* and *fd* statistics were provided, along with the corresponding block jackknife corrected *P* value. **d** Genomic characteristics of the putative introgressed regions between *C.sikamea* and *C. gigas*. Manhattan plot showing the $f_{dM}$ value between *C.sikamea* and *C. gigas*. The red dashed line shows the cutoff value, calculated by highest 1% of $f_{dM}$ values.

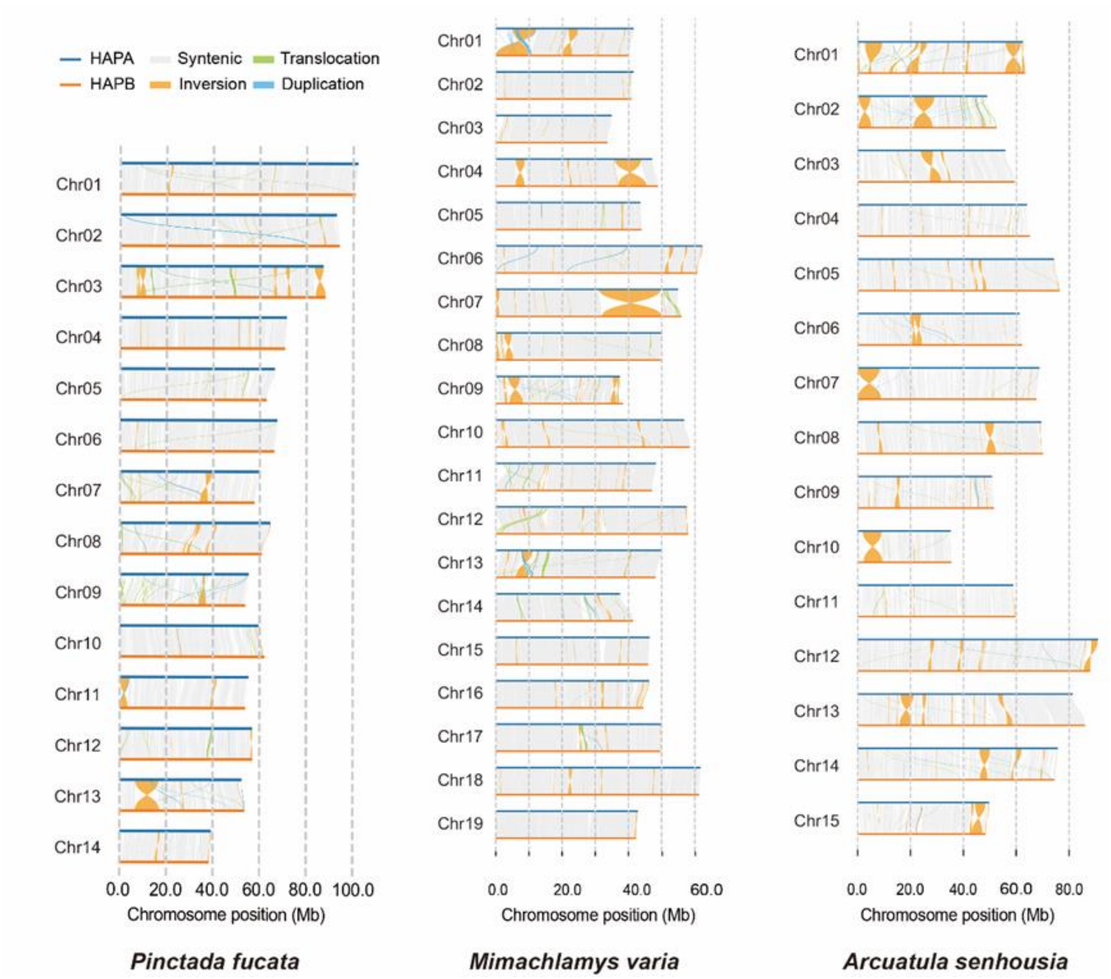


**Supplementary Figure 5. The comparing haplotype genomes of *A. senhousia*, *M. varia*, and *P. fucata*.** Structural rearrangements and synteny are represented by colored lines connecting regions of the haplotype-resolved chromosome-level genome of *A. senhousia, M. varia. a, and P. fucata.*

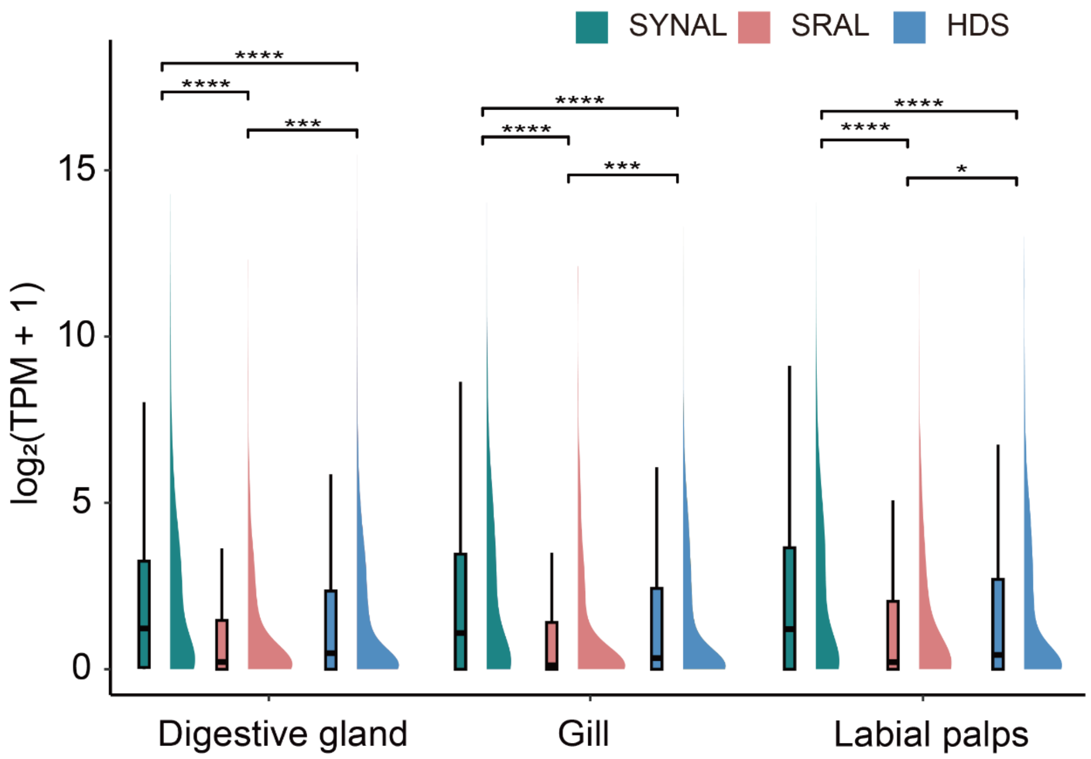


**Supplementary Figure 6.** Comparing gene expression levels within SYNAL, SRAL, and HDS of *C. sikamea* genome in the tissues of the digestive gland, gill, and labial palps. Significance testing was conducted using the Wilcox test.

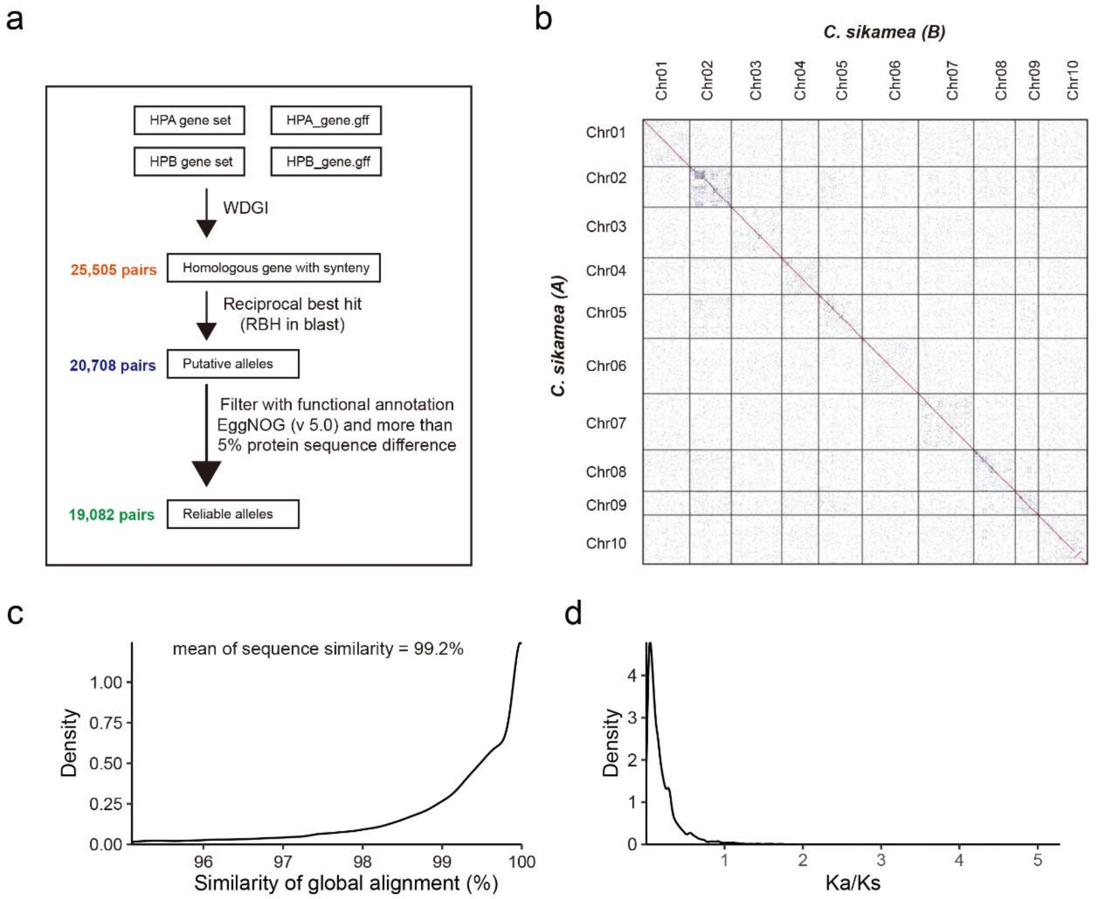


**Supplementary Figure 7. Identified alleles pipeline details and comparative analysis of gene homology between haplotype genomes and sequence comparison of allelic genes. a** Diagram of the approach taken to identify alleles between haplotype genomes. **b** The homologous gene dotplot between haplotype genomes. **c** The sequences similarity of identified alleles. **d** The Ka/Ks ratio of identified alleles.

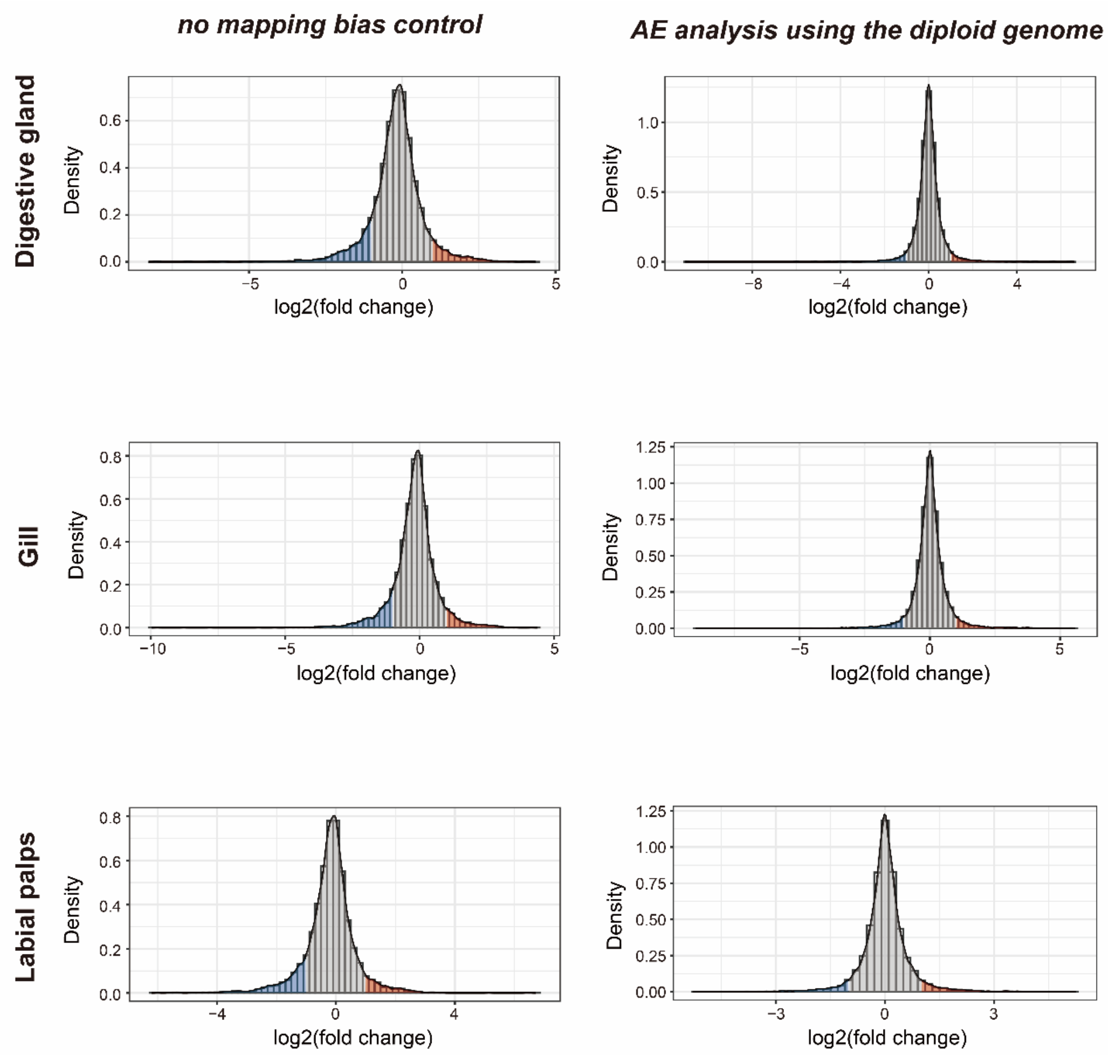


**Supplementary Figure 8. Analysis of the effectiveness of the direct quantification of allele expression (AE) using the diploid genome in mapping bias control.**

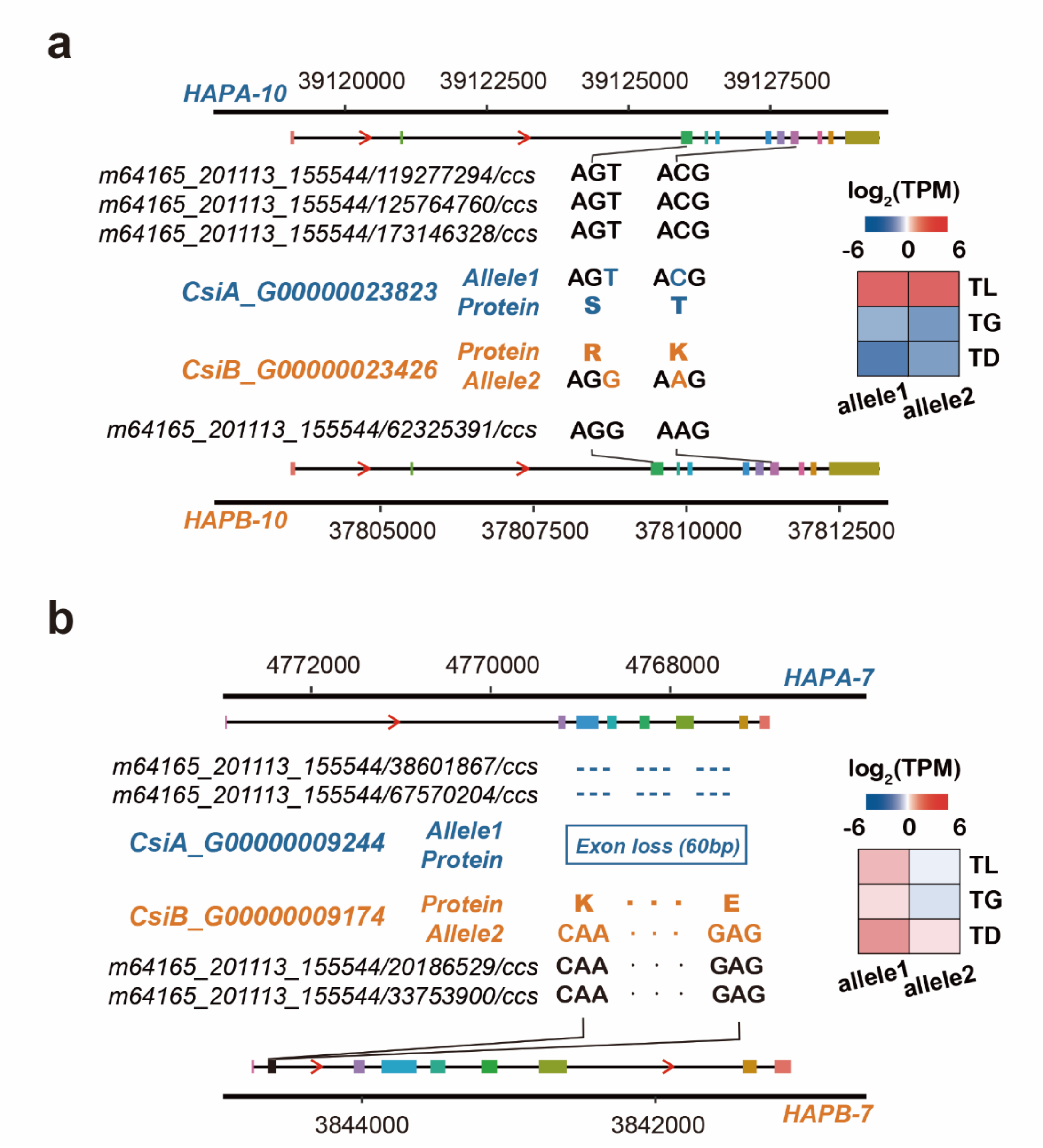


**Supplementary Figure 9. Diagrams of alleles modified by nonsynonymous mutations or structural variations show varying degrees of allele-specific expression (ASE) across different tissues. a** An example of an ASEG with two nonsynonymous mutations, which are supported by Iso-seq reads. Right, allelic differential expression of this gene in three tissues, including the labial palps (TL), gill (TG), and digestive gland (TD). **b** An example of an ASEG with 60-bp deletion, which are supported by Iso-seq reads. Right, allelic differential expression of this gene in three tissues; red arrowheads is gene transcription direction.

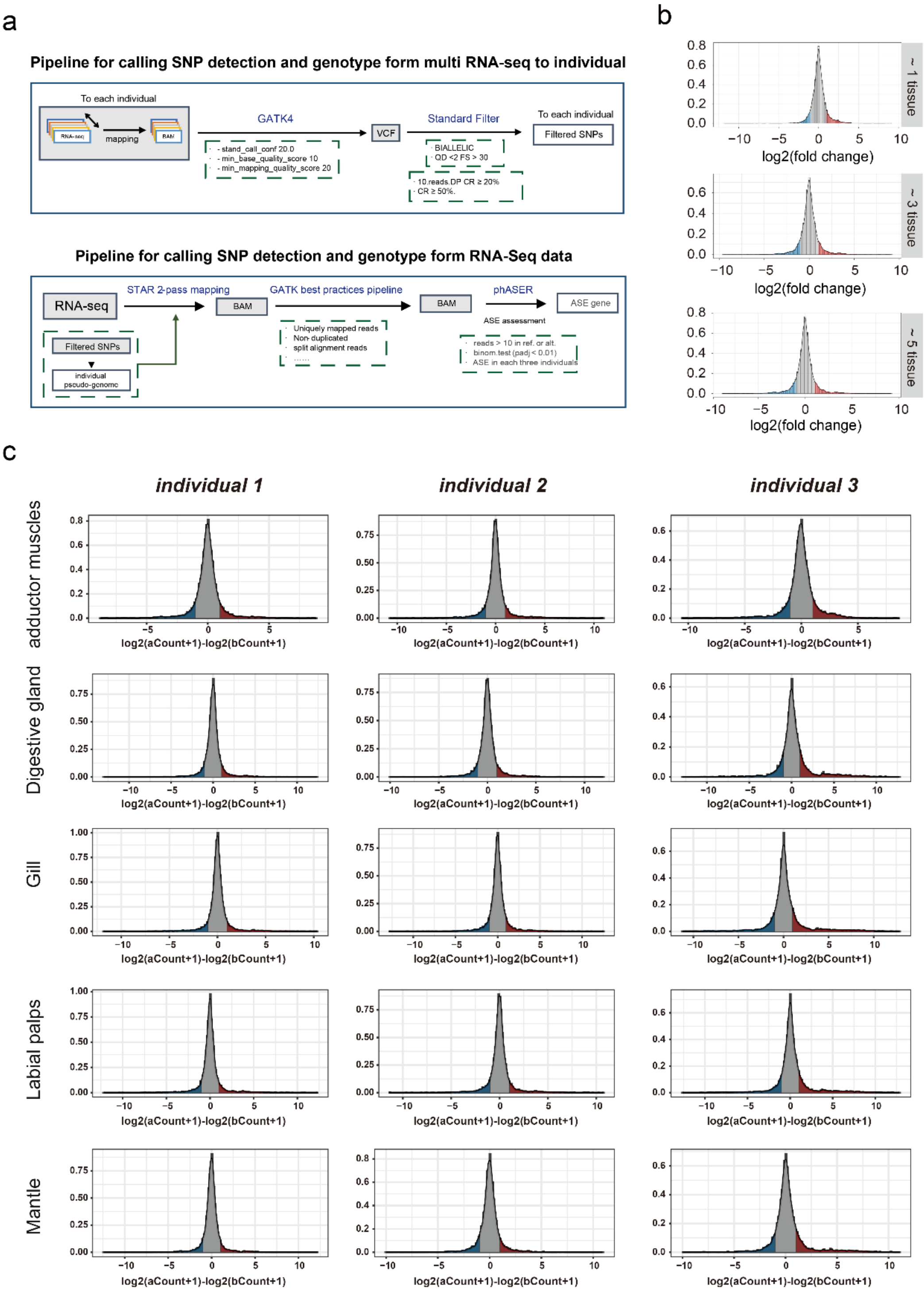


**Supplementary Figure 10. Transcriptome-based SNP detection and ASE analysis process and feasibility analysis. a** Workflow used to detect SNPs and analysis Allele-Specific Expression (ASE) from different RNA-seq data for each individual. **b** With the tissue addition for calling SNP detection, the histograms of a global reference allelic ratio at every gene locus in the adductor muscle. **c** Analysis of the effectiveness of using multi RNA-seq data for individual allele expression (AE) in mapping bias control.

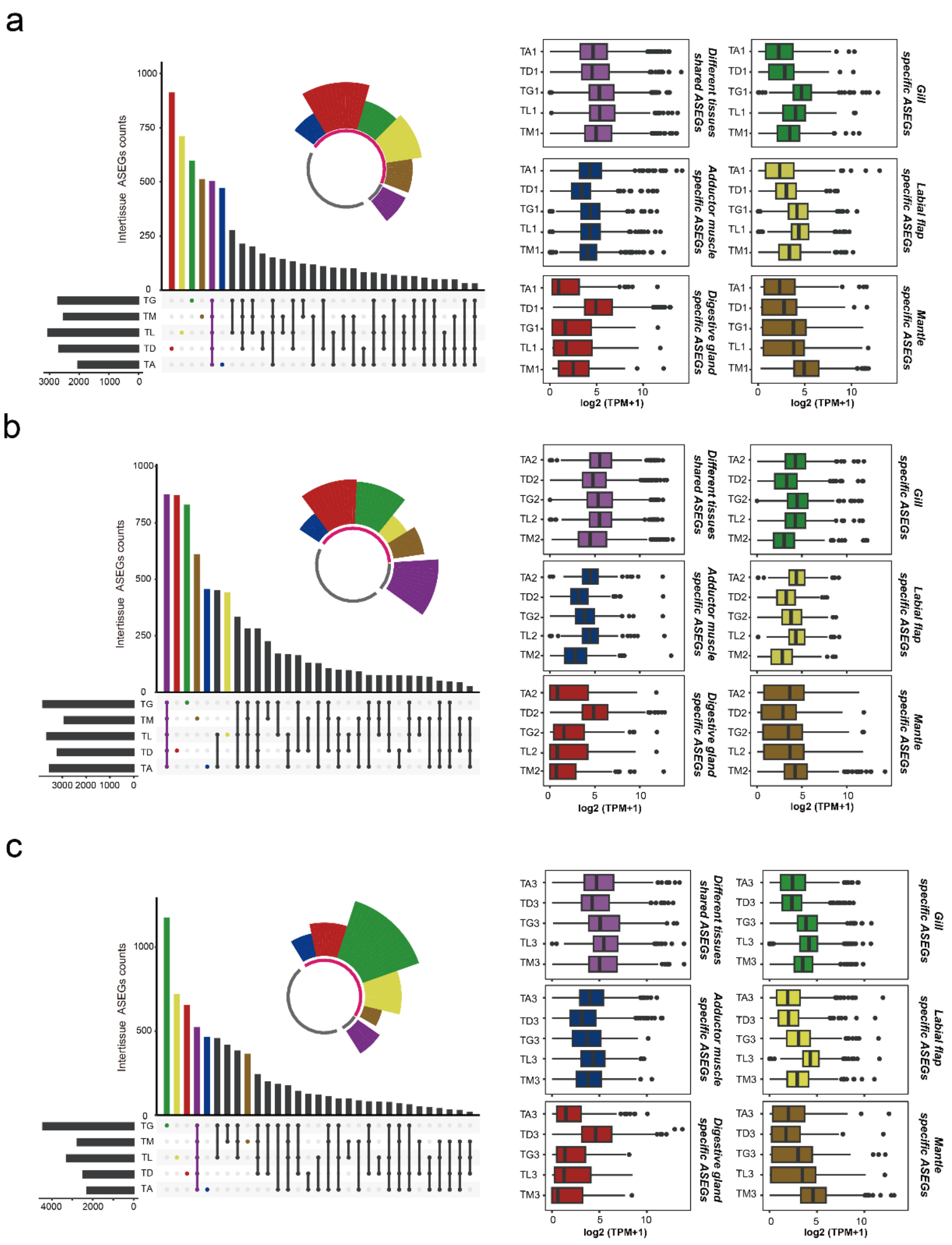


**Supplementary Figure 11. The comparative analysis of allele-specific expression (ASE) gene sets from five different tissues of the *individual 1* (a), *individual 2* (b), and *individual 3* (c).** The upper-right coxcomb chart illustrates the proportion of tissue-specific ASE genes (ASEGs) and tissue-shared ASEGs. The figure on the right compares the overall expression levels of ASEGs shared across all tissues with those of tissue-specific ASEGs in various tissues. The colors represent ASEGs shared across all tissues (purple) and tissue-specific ASEGs for different tissues: adductor muscle (blue), digestive gland (red), gill (green), labial palps (yellow), and mantle (brown).

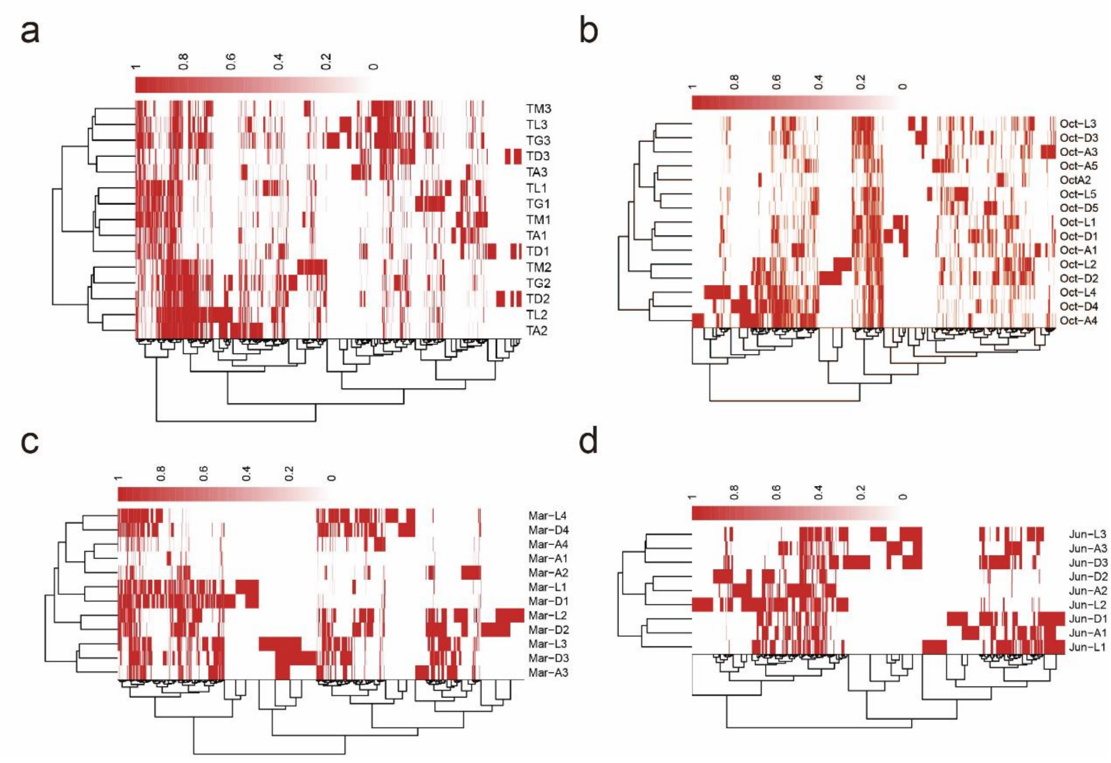


**Supplementary Figure 12. Comparing ASE gene sets from different organizations and tissue-specific ASE gene sets. a** *RNA-seq dataset1* from five different tissues of three individuals (TA, adductor muscles; TD, digestive gland; TG, gill; TL, labial palps; and TM, mantle); **b** *RNA-seq dataset2* from three different tissues of five individuals. **c** *RNA-seq dataset3* from three different tissues of four individuals. **d** *RNA-seq dataset4* from three different tissues of three individuals.

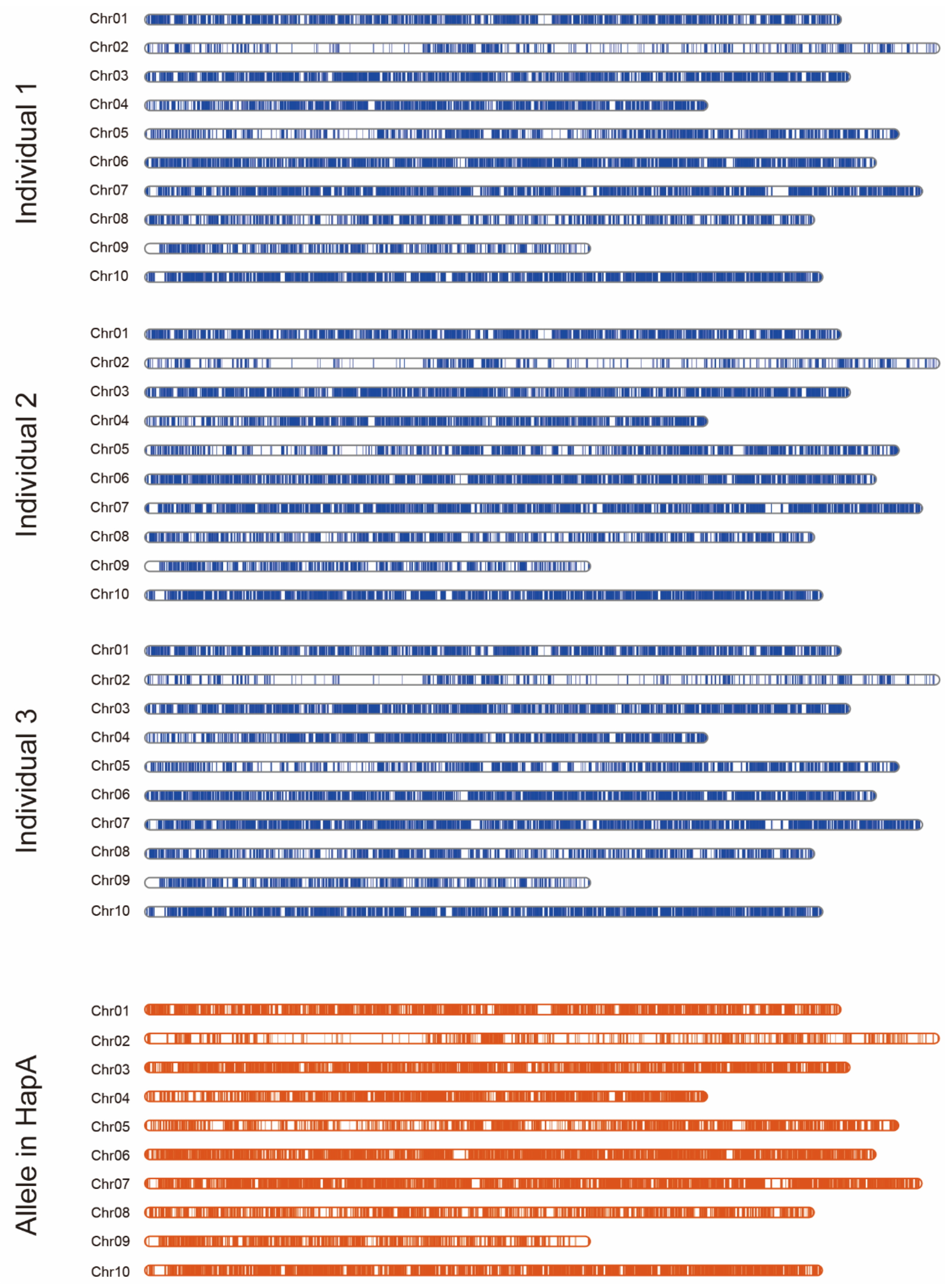


**Supplementary Figure 13. Comparing the distribution of the identified alleles (orange) by haplotype genomes with the distribution of the gene marked by effective SNPs (blue) in each individual.**

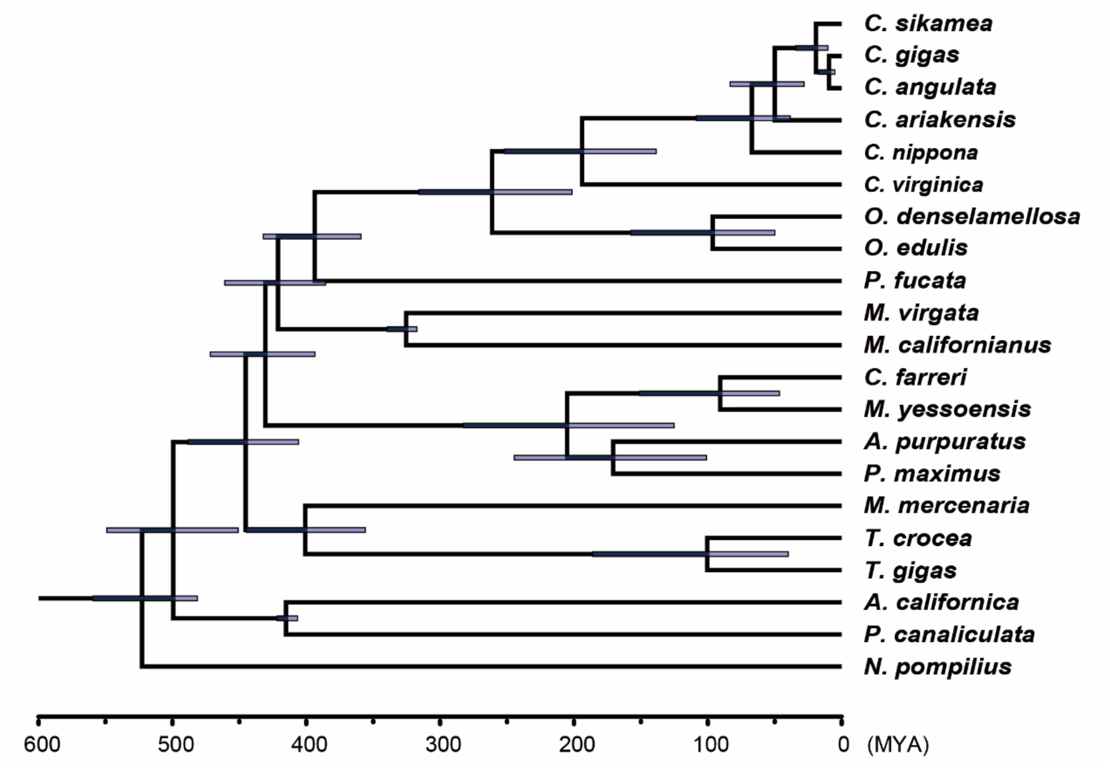


**Supplementary Figure 14. Phylogenetic tree of 21 representative molluscs.**

### Supplementary Tables

#### Supplementary Table 1. Sequencing data for genome assembly of *C. sikamea*

| | |
|---|---|
| Illumina sequencing | |
| Data size of short reads (Gb) | 26.66 (26,655,764,400) |
| Number of short reads | 177,705,096 |
| PacBio-HiFi sequencing | |
| Data size of subreads (Gb) | 440.58Gb (440,578,103,772) |
| Number of subreads | 31,673,400 |
| Data size of CCS reads (Gb) | 29.53 (29,533,700,187.7) |
| Sequencing depth of CCS reads (X) | 57.11 |
| Largest length of CCS reads | 41,426 |
| Average length of CCS reads (bp) | 15,216.38 |
| Number of CCS reads | 1,940,915 (31,673,400) |
| N50 values of CCS reads (bp) | 836,270 |
| Hi-C sequencing | |
| Clean data (Gb) | 50.49 (50,492,423,410 bp) |
| Sequencing depth (X) | 97.63 |
| Iso-Seq sequencing | |
| Data size (Gb) | 54.08 |
| Number of subreads | 24,343,840 |
| Number of CCS reads | 629,581 |
| Number of high-quality FLNC reads | 540,577 |
| Number of consensus reads | 405,659 |

**Supplementary Table 2. Statistics of genome assembly and annotation of *C. sikamea.***

| Feature | *C. sikamea* | |
|---|---|---|
| Estimated genome size (bp) | 517,203,934 | |
| Heterozygosity (%) | 3.31 | |
| Duplication (%) | 0.57 | |
| *De novo* assembly | | |
| Assembly size (Gb) | 1.08 (1,076,379,459) | |
| Contig number | 1,243 | |
| Largest length of contigs | 10,147,741 | |
| Average length of contigs | 865,952.90 | |
| Contig N50 (Mb) | 2,159,660 | |
| BUSCO completeness of assembly (%) | 98.9% | |
| Haplotype-resolved chromosomal-level assembly and annotation | | |
| | Haplotype A | Haplotype B |
| Length of chromosomes (Mb) | 526.67 | 516.10 |
| BUSCO completeness of assembly (%) | 98.2% | 98.0% |
| BUSCO completeness of annotation (%) | 97.8% | 97.1% |
| Number of annotated genes | 30,052 | 29,609 |
| Total number of anchored genes | 59,661 | |

**Supplementary Table 3. Merqury analysis of haplotype-resolved genome.**

| Assembly | solid k-mers in the assembly | Total solid k-mers in the read set | QV | Error Rate (%) | Completeness (%) |
|---|---|---|---|---|---|
| HAPA | 359,743,023 | 525,988,251 | 45.63 | 2.73302e-05 | 68.39 |
| HAPB | 357,744,400 | 525,988,251 | 45.65 | 2.72151e-05 | 68.01 |
| Both | 519,522,144 | 525,988,251 | 45.64 | 2.72733e-05 | 98.77 |

**Supplementary Table 4. Statistical analysis of structural rearrangements and synteny between haplotype-resolved genome.**

| #Structural annotations | *Crassostrea sikamea* | | |
|---|---|---|---|
| Genome Size (reference, bp) | ***Haplotype A*** | | ***526,669,925*** |
| Genome Size (query, bp) | ***Haplotype B*** | | ***516,097,410*** |
| #Variation_type | Count | Length_ref | Length_qry |
| Syntenic regions | 1,968 | 373,596,569 | 384,045,088 |
| Inversions | 115 | 18,259,928 | 19,055,908 |
| Translocations | 2,617 | 9,272,748 | 9,341,588 |
| Duplications (reference) | 203 | 1,197,710 | - |
| Duplications (query) | 685 | - | 2162939 |
| # Haplotype divergent sequences (HDS) | Count | Length_ref | Length_qry |
| Highly divergent region (reference) | 51,953 | 241,616,294 | - |
| Highly divergent region (query) | 51,953 | - | 252,989,386 |
| Un-aligned region (reference) | 4,698 | 124,877,049 | - |
| Un-aligned region (query) | 5,143 | - | 101,499,583 |

**Supplementary Table 5. The average IBD block number and length among four oyster populations.**

| Species | Average IBD block number | Avearge IBD block length, bp |
|---|---|---|
| *C.gigas-C.angulata* | 166.62 | 405877.05 |
| *C.gigas-C. sikamea* | 2.17 | 4347.82 |
| *C.angulata-C. sikamea* | 3.55 | 5728.81 |
| *C. ariakensis - C.gigas* | 0.06 | 132.95 |
| *C. ariakensis - C.angulata* | 0.05 | 148.31 |
| *C. ariakensis- C. sikamea* | 0.18 | 364.81 |

**Supplementary Table 6. Genome assembly statistics of *A. senhousia* and *M. varia*.**

| Feature | *Arcuatula senhousia* | |
|---|---|---|
| *De novo* assembly | | |
| Assembly size (Gb) | 2.05 (2,050,023,122) | |
| Contig number | 4,460 | |
| Largest length of contigs | 14,230,821 | |
| Average length of contigs | 459,646.4 | |
| Contig N50 (bp) | 3,336,361 | |
| BUSCO completeness of assembly (%) | 98.3% | |
| Haplotype-resolved chromosomal-level assembly and annotation | | |
| | Haplotype A | Haplotype B |
| Length of chromosomes (pb) | 948,494,193 | 961,024,312 |
| BUSCO completeness of assembly (%) | 97.6% | C:97.8% |
| Feature | *Mimachlamys varia* | |
| *De novo* assembly | | |
| Assembly size (Gb) | 2.07 (2,073,743,215) | |
| Contig number | 5,959 | |
| Largest length of contigs | 10,206,222 | |
| Average length of contigs | 348,001.9 | |
| Contig N50 (bp) | 1,989,882 | |
| BUSCO completeness of assembly (%) | 99.7% | |
| Haplotype-resolved chromosomal-level assembly and annotation | | |
| | Haplotype A | Haplotype B |
| Length of chromosomes (bp) | 911,008,730 | 909,581,060 |
| BUSCO completeness of assembly (%) | 96.0% | 96.1% |

**Supplementary Table 7. The** Hypergeometric test result of lineage-specific gene in different phylogenetic ‘strata’ within HDS.

| group | Number of Genes in Genome | Number of Genes in HDS | Number of lineage-specific gene in HDS | Number of lineage-specific gene in HDS | p_value | padj |
|---|---|---|---|---|---|---|
| I | 30,052 | 11,131 | 421 | 137 | 0.976660351 | 0.976660351 |
| II | 30,052 | 11,131 | 323 | 120 | 0.504345648 | 0.605214778 |
| III | 30,052 | 11,131 | 1601 | 633 | 0.018104126 | 0.027156188 |
| IV | 30,052 | 11,131 | 332 | 142 | 0.017731944 | 0.027156188 |
| V | 30,052 | 11,131 | 403 | 214 | 2.98E-11 | 8.95E-11 |
| VI-a | 30,052 | 11,131 | 819 | 682 | 8.92E-168 | 5.35E-167 |
| VI-b | 29,936 | 11,686 | 929 | 633 | 3.77E-74 | 1.31E-73 |

### Supplementary Table 8. The KEGG enrichment results of genes in HDS of two haploid genomes

| # The KEGG enrichment results of genes in HDS of HPA genome | | | | | |
|---|---|---|---|---|---|
| **ID** | **Description** | **GeneRatio** | **BgRatio** | **pvalue** | **qvalue** |
| ko03032 | **DNA replication proteins [BR:ko03032]** | 111/3403 | 247/12758 | 3.54E-10 | 1.31E-07 |
| ko04091 | **Lectins [BR:ko04091]** | 164/3403 | 412/12758 | 2.61E-09 | 4.82E-07 |
| ko03029 | **Mitochondrial biogenesis [BR:ko03029]** | 177/3403 | 471/12758 | 8.67E-08 | 1.07E-05 |
| ko04054 | **Pattern recognition receptors [BR:ko04054]** | 136/3403 | 345/12758 | 1.19E-07 | 1.09E-05 |
| ko03011 | **Ribosome [BR:ko03011]** | 77/3403 | 177/12758 | 8.59E-07 | 6.35E-05 |
| ko03010 | **Ribosome** | 63/3403 | 140/12758 | 2.11E-06 | 1.30E-04 |
| ko03430 | **Mismatch repair** | 26/3403 | 44/12758 | 5.82E-06 | 3.07E-04 |
| ko03018 | **RNA degradation** | 57/3403 | 127/12758 | 6.99E-06 | 3.23E-04 |
| ko03440 | **Homologous recombination** | 36/3403 | 72/12758 | 1.96E-05 | 8.07E-04 |
| ko05206 | MicroRNAs in cancer | 137/3403 | 393/12758 | 1.72E-04 | 6.35E-03 |
| ko04621 | **NOD-like receptor signaling pathway** | 96/3403 | 262/12758 | 2.23E-04 | 7.50E-03 |
| ko04514 | Cell adhesion molecules | 80/3403 | 212/12758 | 2.50E-04 | 7.69E-03 |
| ko04217 | Necroptosis | 73/3403 | 199/12758 | 1.15E-03 | 2.74E-02 |
| ko04215 | **Apoptosis - multiple species** | 49/3403 | 124/12758 | 1.19E-03 | 2.74E-02 |
| # The KEGG enrichment results of genes in HDS of HPB genome | | | | | |
| **ID** | **Description** | **GeneRatio** | **BgRatio** | **pvalue** | **qvalue** |
| ko03018 | **RNA degradation** | 68/3307 | 135/12629 | 1.41927E-09 | 5.09443E-07 |
| ko03032 | **DNA replication proteins [BR:ko03032]** | 108/3307 | 258/12629 | 2.41959E-08 | 4.34253E-06 |
| ko04054 | **Pattern recognition receptors [BR:ko04054]** | 127/3307 | 329/12629 | 3.98549E-07 | 4.7686E-05 |
| ko03029 | **Mitochondrial biogenesis [BR:ko03029]** | 173/3307 | 478/12629 | 5.72262E-07 | 5.1353E-05 |
| ko03440 | **Homologous recombination** | 38/3307 | 75/12629 | 4.87834E-06 | 0.000301958 |
| ko03011 | **Ribosome [BR:ko03011]** | 73/3307 | 175/12629 | 5.0474E-06 | 0.000301958 |
| ko04091 | **Lectins [BR:ko04091]** | 132/3307 | 362/12629 | 8.07121E-06 | 0.000413877 |
| ko03010 | **Ribosome** | 59/3307 | 137/12629 | 1.25006E-05 | 0.000560882 |
| ko04215 | **Apoptosis - multiple species** | 52/3307 | 125/12629 | 0.000119709 | 0.003929265 |
| ko04621 | **NOD-like receptor signaling pathway** | 93/3307 | 253/12629 | 0.000120413 | 0.003929265 |
| ko00601 | Glycosphingolipid biosynthesis - lacto and neolacto series | 49/3307 | 119/12629 | 0.000248805 | 0.006869848 |
| ko04217 | Necroptosis | 71/3307 | 190/12629 | 0.000423403 | 0.010855672 |
| ko04624 | Toll and Imd signaling pathway | 62/3307 | 162/12629 | 0.000462224 | 0.011060948 |
| ko03430 | **Mismatch repair** | 20/3307 | 39/12629 | 0.000701787 | 0.015744042 |
| ko04625 | C-type lectin receptor signaling pathway | 88/3307 | 250/12629 | 0.000916929 | 0.018284966 |
| ko04623 | Cytosolic DNA-sensing pathway | 57/3307 | 155/12629 | 0.002273656 | 0.032921182 |

**Supplementary Table 9. Compare the SNP detection from RNA-Seq Data and DNA-Seq Data.**

| # The SNP detection from RNA Seq Data | | | | |
|---|---|---|---|---|
| **Indvidual** | **raw SNP** | **filtered biallelic SNPs** | **filtered biallelic SNPs with same GT in DNA-seq** | **high-quality heterozygous SNPs *** |
| Ind. 1 | 1,863,590 | 841,265 | 765,195 | 434,226 |
| Ind. 2 | 1,786,962 | 849,020 | 764,097 | 430,655 |
| Ind. 3 | 1,595,200 | 806,211 | 722,090 | 408,230 |
| # The SNP detection from DNA-Seq Data | | | | |
| **Indvidual** | **raw SNP** | **filtered biallelic SNPs** | **filtered biallelic SNPs in exon region** | **filtered biallelic SNPs in promoter region** |
| Ind. 1 | 12,899,540 | 11,686,908 | 1,108,354 | 1,386,857 |
| Ind. 2 | 12,567,299 | 11,503,695 | 1,094,716 | 1,363,942 |
| Ind. 3 | 12,956,144 | 11,728,293 | 1,108,727 | 1,396,343 |

* High-quality heterozygous SNPs are defined as those showing the same genotype in at least two tissue types, and if the hetSNP is only identified in one tissue, they are retained upon validation by DNA sequencing data.

**Supplementary Table 10. The Statistical Analysis of gene expression and allele-specific expression (ASE) in *RNA-seq dataset 1*.**

| ***Individual 1*** | **Gene count** | **Expression gene counts (TPM > 0.1)** | **Mark gene counts** | **no ASEG counts (padj>0.1)*** | **ASEG counts (padj<0.05)** |
|---|---|---|---|---|---|
| **Adductor muscles** | 30,052 | 18,660 | 8,022 | 5,624 | 2,034 |
| **Digestive gland** | 30,052 | 21,141 | 10,337 | 7,169 | 2,667 |
| **Gill** | 30,052 | 20,890 | 10,307 | 7,066 | 2,711 |
| **Labial palps** | 30,052 | 20,926 | 10,345 | 6,741 | 3,049 |
| **Mantle** | 30,052 | 20,966 | 10,182 | 7,090 | 2,519 |
| **ALL** | 30,052 | 25,065 | 12,194 | 5,718 | 6,390 |
| ***Individual 2*** | **Gene count** | **Expression gene counts (TPM > 0.1)** | **Mark gene counts** | **no ASEG counts (padj>0.1)*** | **ASEG counts (padj<0.05)** |
| **Adductor muscles** | 30,052 | 21,251 | 11,117 | 6,937 | 3,530 |
| **Digestive gland** | 30,052 | 20,677 | 11,059 | 7,330 | 3,199 |
| **Gill** | 30,052 | 20,769 | 11,052 | 6,624 | 3,810 |
| **Labial palps** | 30,052 | 21,274 | 11,191 | 6,855 | 3,647 |
| **Mantle** | 30,052 | 19,914 | 10,370 | 6,894 | 2,921 |
| **ALL** | 30,052 | 25,061 | 13,064 | 5,470 | 7,526 |
| ***Individual 3*** | **Gene count** | **Expression gene counts (TPM > 0.1)** | **Mark gene counts** | **no ASEG counts (padj>0.1)*** | **ASEG counts (padj<0.05)** |
| **Adductor muscles** | 30,052 | 19,526 | 9,176 | 6,439 | 2,280 |
| **Digestive gland** | 30,052 | 19,241 | 9,719 | 6,776 | 2,467 |
| **Gill** | 30,052 | 20,021 | 11,086 | 6,031 | 4,427 |
| **Labial palps** | 30,052 | 20,982 | 11,002 | 7,087 | 3,278 |
| **Mantle** | 30,052 | 20,181 | 10,441 | 7,138 | 2,751 |
| **ALL** | 30,052 | 24,504 | 12,758 | 5,306 | 7,386 |

* Statistical analysis of genes with no allele-specific expression (ASE) across all tissues in individuals

**Supplementary Notes**

**Supplementary Note 1**

**Fluctuations of effective population size (Ne) for each oyster species**

Whole-genome resequencing of *C. sikamea* from a wild population (Ningde, China), in addition with publicly available datasets generated from the other three widely distributed oyster species, including *C. ariakensis* (n=28), *C. angulata* (n=21), and *C. gigas* (n=21) were used in this study (Fig. S3a; Supplementary Data 10). Analyses of the population structure in these four closely related *Crassostrea* oyster species revealed the highest nucleotide diversity in the *C. sikamea* (Π=0.0585) (Fig. S3b). The decay of linkage disequilibrium (LD) with physical distance between SNPs occurs at only 35 bp in *C. sikamea* (decaying to $r^2$ of 0.2), whereas the equivalent distances are 60-580 bp in other oyster species (Fig. S3c). We also assessed fluctuations of effective population size (*Ne*) for each oyster species with the pairwise sequentially Markovian coalescent method. Over the past million years, the impact of glaciation events on the four oyster species has varied greatly. The *Ne* of *C. gigas* and *C. ariakensis* reached its peak before the Mindel glaciation (MG, 0.68-0.80 Mya) (Fig. S3d). In contrast, the *Ne* of *C. angulata* peaked after the MG, while *C. sikamea* reached their first peak just before the Riss glaciation (RG, 0.24-0.37 Mya). During the RG and Würm glaciation (WG, 0.01-0.12 Mya) periods, the Ne of all four oyster species showed a sustained decline. At the end of the last glaciation period (WG), the Ne of all species reached their bottom, while the recovered Ne of *C. sikamea* was significantly higher than that of other oyster species.

We find no evidence of gene flow derived from recent interspecific hybridization among these *Crassostrea* oyster species with the ADMIXTURE analysis, even for closely related species with the same domain distribution (Fig. S4a). Detection of shared identical-by-descent (IBD) haplotypes within and between oyster species showed that *C. sikamea* shared a few IBD haplotypes with *C. angulata* and *C. gigas* (Fig. S4b), suggesting ancient interspecific introgression. We further verify the existence of this ancient introgression by D-statistics and *fd*, which revealed a closer genetic relationship between *C. sikamea* and *C. gigas* than with *C. angulata* (Fig. S4c). The small *D* and *fd* values may be attributed to the occurrence of hybridization between *C. sikamea* and *C. gigas* after speciation, followed by backcrossing and genetic drift, which could have diminished most of the introgression signals. Limited introgression suggests that hybridization has a minimal impact on the genomic characteristics and genetic structure of oysters in nature.

As we find ancient gene flow between *C. sikamea* and geographically distant *C. gigas*, although its impact on shaping the complex genome of oysters may be minimal. Considering the dispersal of East Asian *Crassostrea* species from the northwest Pacific to the tropical western Pacific regions, along with the observed shared distribution of *C. gigas* and *C. sikamea* in the waters of Korea and Japan[1,2], we hypothesize that the initial differentiation of East Asian *Crassostrea* species occurred in the northwest Pacific. It is during this period that gene flow events between *C. sikamea* and *C. gigas* likely took place. We implemented pairwise sequentially Markovian coalescent (PSMC)[3] to estimate effective population size (*Ne*) dynamics over the past several million years for four oyster species. Three *C. sikamea* individuals, three *C. gigas* individuals, two *C. angulata* individuals, and two *C .ariakensis* individuals were included in the PSMC analysis. The parameters were set as follows: "*-N25 -t15 -r5 -p "4+25*2+4+6"*". The generation time (*g*) was set to 1, and mutation rate (μ) to $0.2 \times 10^{-8}$ for all four species (Fig. S4d). Before the MG period, with notable differences in the fluctuations of effective population size (*Ne*) compared to other *Crassostrea* species, it is possible that *C. sikamea* expanded and settled in the coastal regions of southern China earlier. The surge in genetic diversity is commonly viewed as a byproduct of historical population expansion[4]. Considering the swift differentiation in the genetic background of *C. sikamea* across various regions[5], the significant expansion of the *C. sikamea* population is expected to contribute to its relatively higher genetic diversity and heterozygosity.

**Supplementary Note 2**

**Comparative analysis of SNPs detected by RNASeq and DNASeq**

Using the RNA-seq data filtered with the filters suggested by GATK, we found 806,211~ 849,020 biallelic SNPs for each individual. We also used high-depth DNA-seq data (~20×) obtained from the mantle for each individual to detect SNPs. With the standard criteria of GATK, we identified 11,503,695 ~ 11,728,293 biallelic SNPs from each individual and then provided a comparison of the SNPs detected from RNA-seq data (Supplementary Table S9). As we can only capture the variant information in the region of expressed genes from the RNA-seq data, the number of SNPs detected by RNA-seq is much lower than that detected by DNA-seq. However, the SNPs detected by RNA-seq and shared with DNA-seq data demonstrate reasonable precision (~ 90%, Supplementary Table S9), thereby ensuring the reliability of SNP detection by RNA-seq in expressed regions. With further filter, we selected a set of high-quality heterozygous SNPs for each individual (Methods; Supplementary Table S9) and used them for haplotype phase and allele-specific expression (ASE) analysis in each tissue. Genes marked by heterozygous SNPs exhibit a distribution across the genome similar to the alleles identified through the diploid genome (Fig. S13).

**Supplementary Note 3**

**Gene family cluster and phylogenetic analysis**

Annotated protein sets of 21 Mollusca representative species were selected for gene family cluster analysis. Among these, the gene sets of eight species within the oyster lineage were reannotated using the same annotation process. And the gene sets of other species were retrieved from NCBI and Ensembl databases. The longest protein isoform was selected to represent each gene. To identify orthogroups and orthologs from the datasets, we followed the OrthoFinder (v2.5.4)[6] pipeline by invoking DIAMOND and OrthoMCL to call orthogroups based on sequence identity. Finally, the gene set was clustered into 36,248 gene families, and a total of 1,184 single-copy gene among these species were identified.

To reveal phylogenetic relationships among oyster lineage species, the CDS sequences of each single-copy orthologous group from the orthology analysis were aligned using MACSE[7]. The alignments were then concatenated into a super-gene alignment after trimming by trimAL[8]. Using the 4-fold degenerate (4D) sites selected from the combined datasets, we generated a Maximum Likelihood (ML) phylogenetic tree using RAxML-NG[9]. The analysis included 200 bootstrap replicates and employed the TIM2 + I + G4 model, as determined by ModelTest-NG[10]. To estimate divergence times, the MCMCtree program in PAML[11] was used for approximate likelihood calculations, based on known approximate divergence times of *A. californica* - *P. canaliculata* (406.1-421.5 Ma), *M. virgata* - *M. californianus* (83.0–103.8 Ma) and a time for the root (480-559.4 Ma) (http://www.timetree.org/).

Molecular clock analysis based on the secondary calibrations suggested that the common ancestor of *C. sikamea* diverged from *C. gigas* and *C. angulata* at 19.17 million years ago (Mya) (10.22-34.25 Mya). Our data support that the divergence of the *Crassostrea* and *Ostrea* genera occurred around 261.12 Mya (201.5-315.9 Mya), which is consistent with previous studies (Fig. S14).
